# Directed Biosynthesis of Mitragynine Stereoisomers

**DOI:** 10.1101/2022.12.22.521574

**Authors:** Carsten Schotte, Yindi Jiang, Dagny Grzech, Thu-Thuy T. Dang, Larissa Laforest, Francisco León, Marco Mottinelli, Satya S. Nadakuduti, Christopher R. McCurdy, Sarah E. O’Connor

**Author notes:** **Corresponding Author**, **Sarah E. O’Connor** – Department of Natural Product Biosynthesis, Max Planck Institute for Chemical Ecology, Hans-Knöll-Straße 8, 07745 Jena Germany;. **Author Contributions**, These authors contributed equally (Carsten Schotte, Yindi Jiang).

## Abstract

*Mitragyna speciosa* (“Kratom”) is used as a natural remedy for pain and management of opioid dependence. The pharmacological properties of Kratom have been linked to a complex mixture of monoterpene indole alkaloids, most notably mitragynine. Here, we report the central biosynthetic steps responsible for the scaffold formation of mitragynine and related corynanthe-type alkaloids. We illuminate the mechanistic basis by which the key stereogenic centre of this scaffold is formed. These discoveries were leveraged for the enzymatic production of mitragynine, the C-20 epimer speciogynine, and a series of fluorinated analogues.

---

*Mitragyna speciosa* (“Kratom”) is an evergreen tree of the *Rubiaceae* family. Kratom consumption leads to stimulating effects at lower doses and opioid-like effects at higher doses.^1^ Manual workers have thus used it for centuries to endure heat, increase physical endurance and combat fatigue.^2,3^ Kratom is also consumed for the (self-)treatment of pain, to mitigate opioid withdrawal symptoms and to treat clinical depression; however, rigorous scientific studies that clinically demonstrate Kratom’s therapeutic efficacy are still lacking.^4^ Because of its purported analgesic properties, as well as for a variety of recreational purposes, Kratom is increasingly used worldwide and is regularly consumed by millions of people in the United States alone.^5,6^

The pharmacological effects of Kratom have been linked to a mixture of >50 corynanthe- and oxindole-type alkaloids (Fig. 1a,b).^7^ Most notable among these are the corynanthe-type alkaloid mitragynine (**1**) and the hydroxylated derivative 7OH-mitragynine (**2**). Both **1** and **2** are nanomolar partial agonists at the human *μ*-opioid receptor (hMOR), and **2** was found to be ~10-fold more potent than morphine in tailflick and hot-plate tests in mice.^8,9^ Intriguingly, speciogynine (**3**), the C-20 epimer of mitragynine (**1)**, does not display agonist activity towards hMOR, though speciogynine (**3**), unlike mitragynine (**1**), is a smooth muscle relaxant. These differential bioactivities highlight the importance of the C-20 stereochemistry in the pharmacology of Kratom alkaloids.^10^

**Figure 1.**
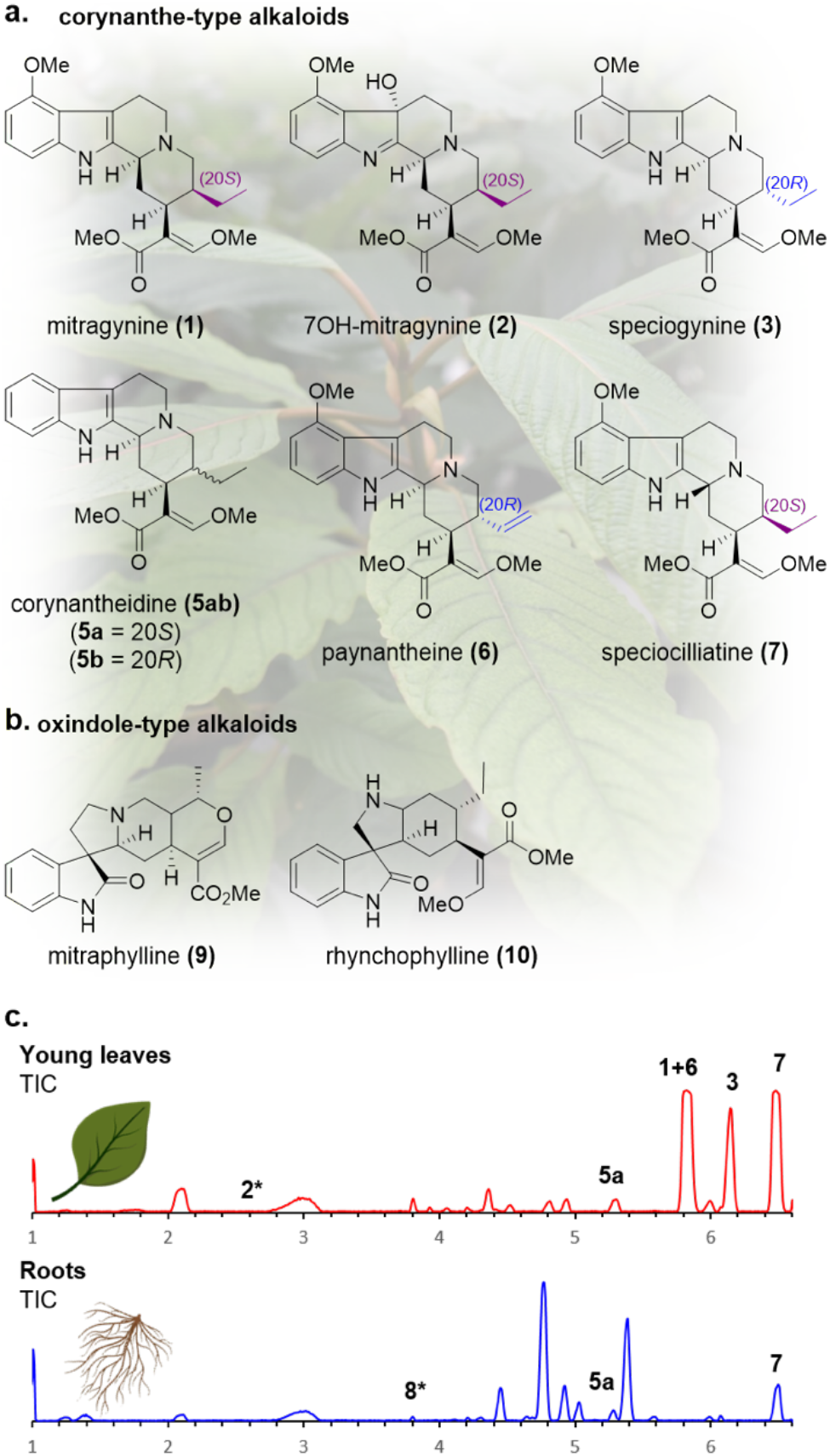
**(a,b)** representative Kratom alkaloids; **(c)** TIC of Kratom leaf and root extracts; * only observed in EIC.

In the present work, we leverage a multi-omics approach to elucidate the key biosynthetic steps that form the corynanthe-type scaffold of Kratom alkaloids. We report the discovery of two medium-chain alcohol dehydrogenases (*Ms*DCS1 and *Ms*DCS2) along with an enol *O*-methyltransferase (*Ms*EnolMT) that converts strictosidine aglycone (**4**) to either of the key stereoisomers (20*S*)-corynantheidine (**5a**) (the precursor to **1**) or (20*R*)-corynantheidine (**5b**) (the precursor to **3**). Rational mutagenesis of MsDCS1 revealed key amino acid residues that control the stereoselective reduction at C-20. A precursor directed biosynthesis approach was then used for the stereoselective production of **1** and **3**, as well as fluorinated analogues.

We first identified where these alkaloids accumulate *in planta* by analyzing methanolic extracts of *M. speciosa* root, stem, bark and leaf tissue at a variety of developmental stages using targeted metabolomics (Fig. 1c; Supplementary Fig. S1-S6). Consistent with literature reports^11,12^ the alkaloid content in the leaves was higher than other organs, with mitragynine (**1**), paynantheine (**6**), speciogynine (**3**) and speciocilliatine (**7**) being the dominant products. Low levels of 7OH-mitragynine (**2**), (20*S*)-corynantheidine (**5a**) and strictosidine (**8**) were also observed. Stem and bark showed similar metabolic profiles, with **7** as the dominant alkaloid and low quantities of **1** and **3** also observed. Notably, root tissue was completely lacking in **1** and **3**, with only **7**, **8** and **5a** detected.

Strictosidine aglycone (**4**) is the central intermediate for most monoterpene indole alkaloids, including **1**, **3** and other Kratom-derived alkaloids. The complete biosynthetic pathway for **4** has been elucidated in the closely related plant *Catharanthus roseus*,^13^ and we readily identified homologues of these biosynthetic genes in the Kratom transcriptome (Scheme 2a). Notably, although **1** and **3** accumulate primarily in leaf and stem, the strictosidine aglycone (**4**) biosynthetic genes were preferentially expressed in roots, suggesting that this organ is the primary site of biosynthesis for the early pathway steps towards **1**. Therefore, either **1** and **3** are produced in root and then subsequently transported to leaf/stem, or alternatively, a biosynthetic intermediate of **1** and **3** is transported to the leaf/stem where the final biosynthetic steps would take place.

Strictosidine aglycone (**4**) is a reactive intermediate that can be reductively trapped into numerous isomers.^14,15^ One isomer, dihydrocorynantheine (**11ab**), has the same scaffold as **1** and **3**, and is therefore a likely biosynthetic intermediate for these alkaloids. Recently, we reported the discovery of a medium-chain alcohol dehydrogenase from *Cinchona pubescens*, dihydrocorynantheine synthase (*Cp*DCS), that converts strictosidine aglycone (**4**) to (20*R*)-dihydrocorynantheine (**11b**) during the biosynthesis of quinine (**12**) (Scheme 1a).^16^ The structure of **11b** was inferred based on (HR)MS/MS experiments and by NMR characterization of the decarboxylated product (20*R*)-dihydrocorynantheal (Supplementary Figure S7).^16^ It seemed logical that orthologous enzymes should catalyze the reduction of strictosidine aglycone to the dihydrocorynantheine scaffold in Kratom (Scheme 1b).^16,17^ Moreover, given the presence of dihy-drocorynantheine-like alkaloids with both (20*S*)- and (20*R*)-stereochemistry in Kratom, we further hypothesized that Kratom would have multiple DCS orthologues with differing stereoselectivity.

**Scheme 1.**
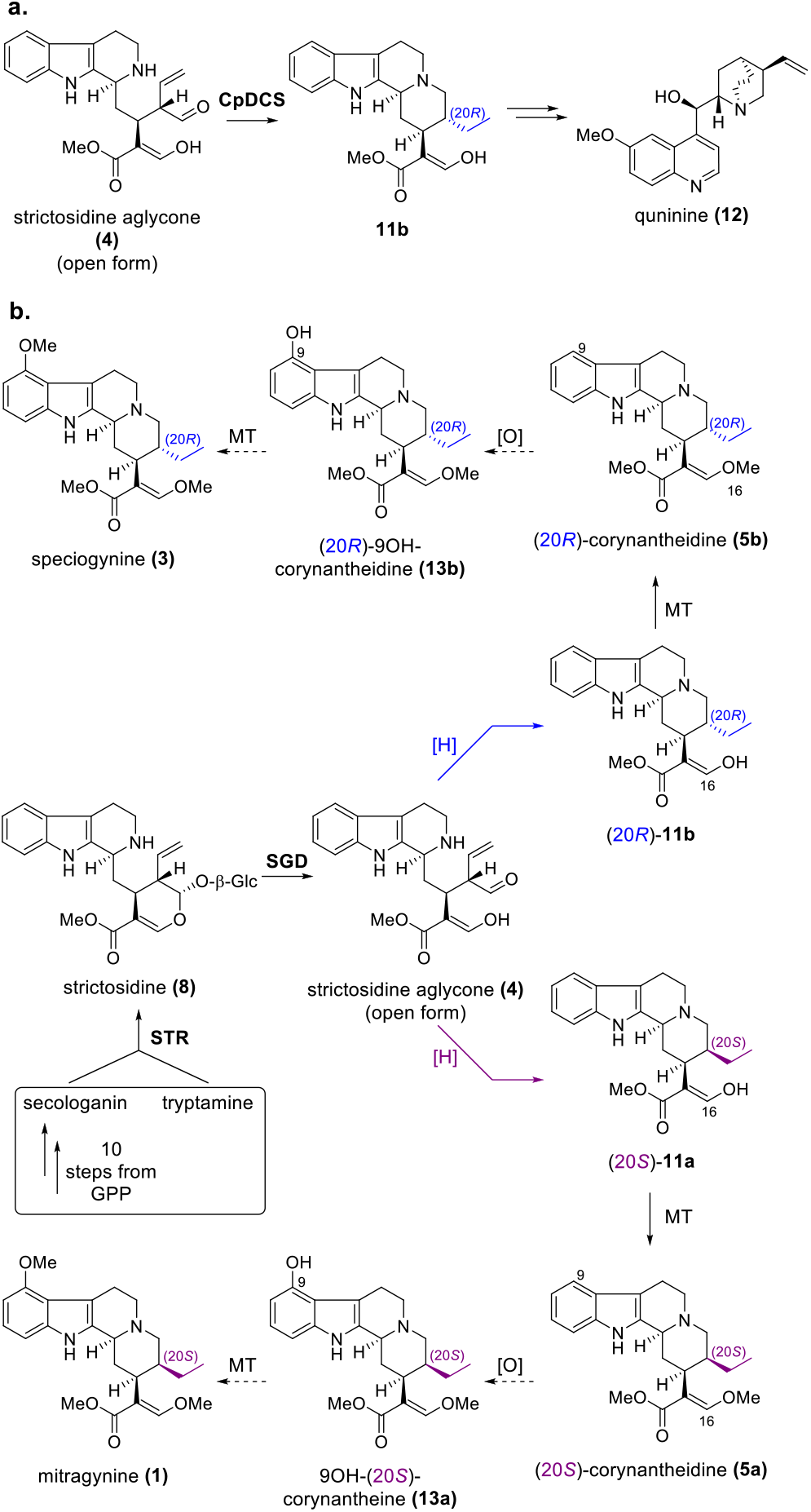
**(a)** Reduction of **4** by *Cp*DCS; **(b**) proposed pathway towards major Kratom alkaloids.

**Scheme 2.**
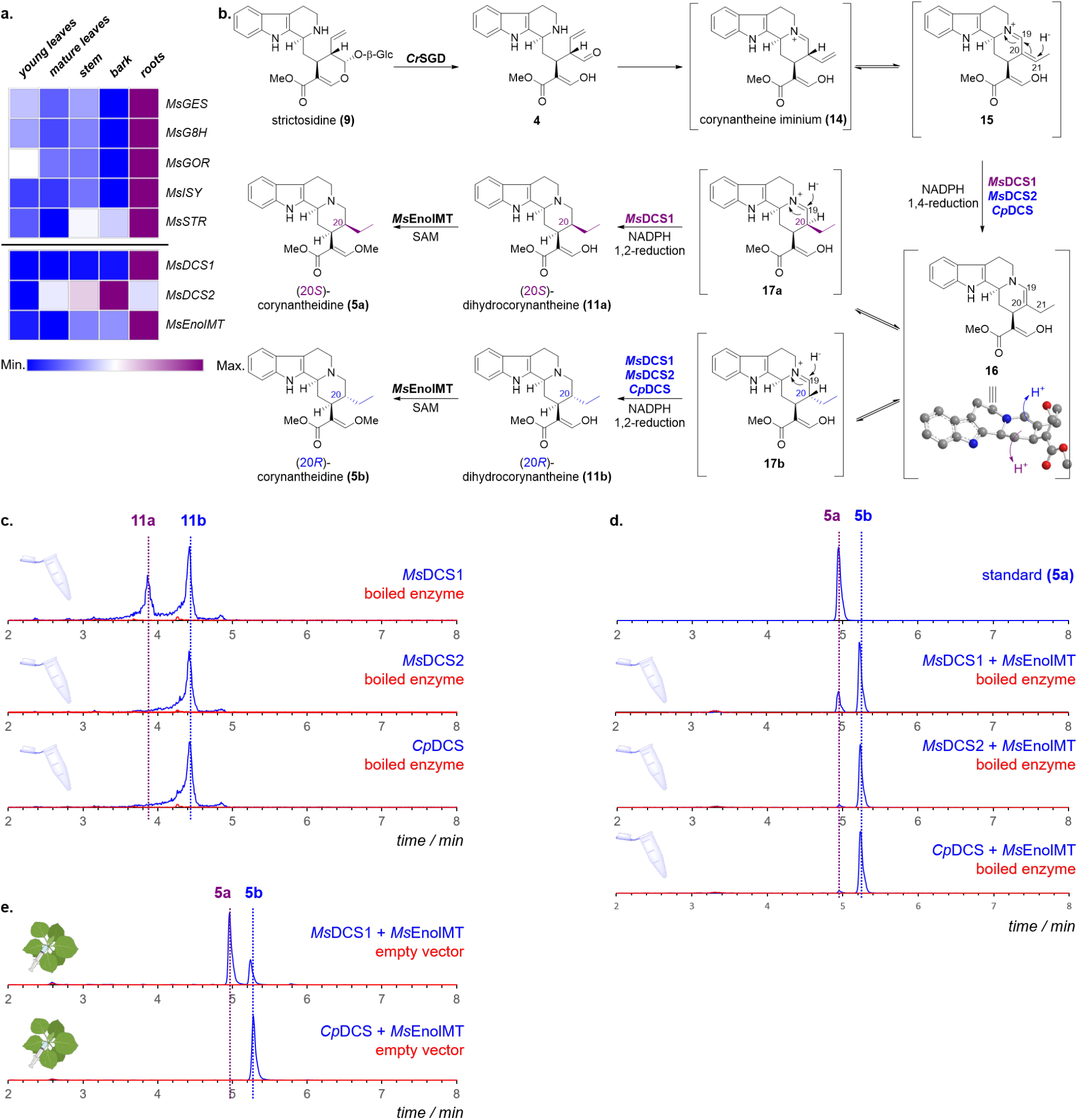
**(a)** Expression profiles of identified genes in Kratom; gene expression levels are represented as FPKM of the *M. speciosa* transcriptome; **(b)** Proposed mechanism for the formation of the corynanthe-type skeleton; **(c)** EIC (*m/z* = 355) of enzymatic assays featuring combinations of **(8)**, *Cr*SGD and *Ms*DCS1/*Ms*DCS2/*Cp*DCS; **(d)** EIC (*m/z* = 369) of enzymatic assays featuring combinations of **(8)**, *Cr*SGD, *Ms*EnolMT and *Ms*DCS1/*Ms*DCS2/*Cp*DCS; **(e)** EIC (*m/z* 369) corresponding to transient expression of *Cr*STR, *Cr*SGD, *Ms*DCS1/*Cp*DCS and *Ms*EnolMT in *Nicotiana benthamiana*.

To identify enzyme candidates from Kratom that catalyze formation of **11a** or **11b** from strictosidine aglycone (**4**), we used the protein sequence of *Cp*DCS to mine the Kratom transcriptome. From this process, 27 gene candidates were expressed in *E. coli*. To assay for enzymatic activity, strictosidine (**8**) was deglycosylated *in situ* with strictosidine glucosidase from *C. roseus* (*Cr*SGD) and incubated with Kratom reductase candidates and NADPH. *Cp*DCS, which afforded (20*R*)-dihydrocorynantheine (**11b**),^16^ was used as a positive control (Scheme 2c; T_R_ = 4.4 min; [M+H]^+^ calculated C_21_H_27_N_2_O_3_ 355.2022, found 355.2014).^16^ Two of the tested Kratom candidates, *Ms*DCS1 and *Ms*DCS2, also produced **11b** (Scheme 2c; Supplementary Fig. S8). Intriguingly, *Ms*DCS1 also yielded a second product with the same HRMS ([M+H]^+^ calculated C_21_H_27_N_2_O_3_ 355.2022, found 355.2015) and MSMS fragmentation pattern as **11b**, but a different retention time (T_R_ = 3.9 min; Scheme 2c; Supplementary Fig. S8). We assumed that this product was (20*S*)-dihydrocorynantheine (**11a**), but due to poor stability, this compound could not be characterized.

Subsequent *O*-methylation at C-16 of **11ab** would yield corynantheidine (**5ab**), which is the next predicted intermediate in the biosynthesis of **1** and **3**. To identify gene candidates that catalyze *O*-methylation at C-16 of **11ab**, we identified annotated methyltransferase genes that coexpress with *Ms*DCS1 (*r* > 0.8, Pearson correlation coefficient). Five purified enzyme candidates were assayed *in vitro* with strictosidine (**8**), strictosidine glucosidase (*Cr*SGD), NADPH, SAM and either *Cp*DCS, *Ms*DCS1 or *Ms*DCS2. A single enzyme, *Ms*EnolMT (*r* = 0.96), showed methyltransferase activity in all enzyme assays (Scheme 2d). Co-incubation of *Ms*EnolMT with *Ms*DCS1 afforded two products with the expected nominal mass of 368 corresponding to the methylated product of **11ab** (HRMS: [M+H]^+^ calculated C_22_H_28_N_2_O_3_ 369.2178, found 369.2164 and 369.2167). The minor product (T_R_ = 4.9 min) eluted at the same retention time and displayed identical MSMS spectra compared to an authentic standard of **5a** (Supplementary Fig. S9),^18^ validating that *Ms*DCS1 generates (20*S*)-dihydrocorynantheine (**11a**). The major product **5b** (TR = 5.4 min) displayed identical MSMS patterns to **5a** (Supplementary Fig. S9, but different retention time. This product has the same retention time and MSMS pattern as the product of CpDCS, which had been previously established to have *R* stereoselectivity (Scheme 2c,d).^18^ Moreover, the MSMS and the retention time of the decarboxylated major MsDCS1 product matched an authentic standard of (20*R*)-dihydrocorynantheal (Supplementary Fig. 7).^18^ *Ms*DCS1 also showed R stereoselectivity (Scheme 2c.

The ratios of **11a** and **11b** produced by *Ms*DCS1 varied considerably among *in vitro* assays, suggesting that assay conditions affect the stereochemical outcome. Therefore, to further corroborate *Ms*DCS1 activity, we transiently expressed *Ms*DCS1 together with strictosidine synthase (*Cr*STR), *Cr*SGD and *Ms*EnolMT in *Nicotiana benthamiana* leaves. Infiltration with tryptamine and secologanin (**19**) (the biosynthetic precursors to strictosidine (**8**)) afforded reproducible ratios of **5a** and **5b**, with **5a** as the dominant product (Scheme 2e). Exchange of *Ms*DCS1 with *Cp*DCS afforded solely **5b**, in agreement with previous observations.

Formation of **11ab** may likely proceed via an initial 1,4-reduction of the α,β-unsaturated iminium dehydrogeissoschizine (**15**), which can form *in situ* upon deglycosidation of **8** (Scheme 2b).^16^ The reduced intermediate **16** tautomerizes to the iminium form, **17ab**, upon protonation at C-20, after which a second 1,2-reduction would occur at C-19. Protonation at C-20 during tautomerization would therefore define the stereochemical outcome. We hypothesized that differences within the active site of these enzymes controlled the face of protonation (*vide infra*).

To identify candidate amino acid residues that direct the C-20 stereochemistry, we generated a structural model of *Ms*DCS1 based on similar reductase enzymes (Fig. 2a-c).^19,20^ Seven amino acids in the binding pocket differentiate *Ms*DCS1 from *Ms*DCS2/*Cp*DCS (Fig. 2c,d; Supplementary Fig. S10). These residues from *Ms*DCS2/*Cp*DCS were introduced into *Ms*DCS1 to swap stereoselectivity at C-20. Assays were performed by transient expression of the resulting mutants in *N. benthamiana* leaves (together with *Cr*STR, *Cr*SGD, *Ms*EnolMT, tryptamine and secologanin; Fig. 2e; Supplementary Figure S11). Mutagenesis of residues 295-298 (SGAS to ATGG) was sufficient to invert the ratio between **5a** and **5b** (from ~74% **5a** in wild-type *Ms*DCS1 to < 35% **5a** in the mutant). In a septuple mutant of *Ms*DCS1 (T53F, I100M, N116S, SGAS295-298ATGG) formation of the (20*S*)-isomer **5a** was nearly abolished (< 5 %). Similar results were observed in analogous mutations in the background of *Cp*DCS, which forms the (20*R*) product **5b** (Fig. 2f). In the septuple *Cp*DCS mutant (F53T, M100I, S116N, ATGG295-298SGAS) the amount of **5a** changed to ~45 % in the mutant compared to 0% in the wild type enzyme (Fig. 2f; compare Supplementary Figure S11 for additional mutations and analysis). Mining of the Kratom genome revealed that *Ms*DCS1 is the only homologue harboring these residues at these positions, so we speculate that *Ms*DCS1 is solely responsible for production of corynanthe-type alkaloids with (20*S*)-stereochemistry in Kratom (Supplementary Fig. S12).

**Figure 2.**
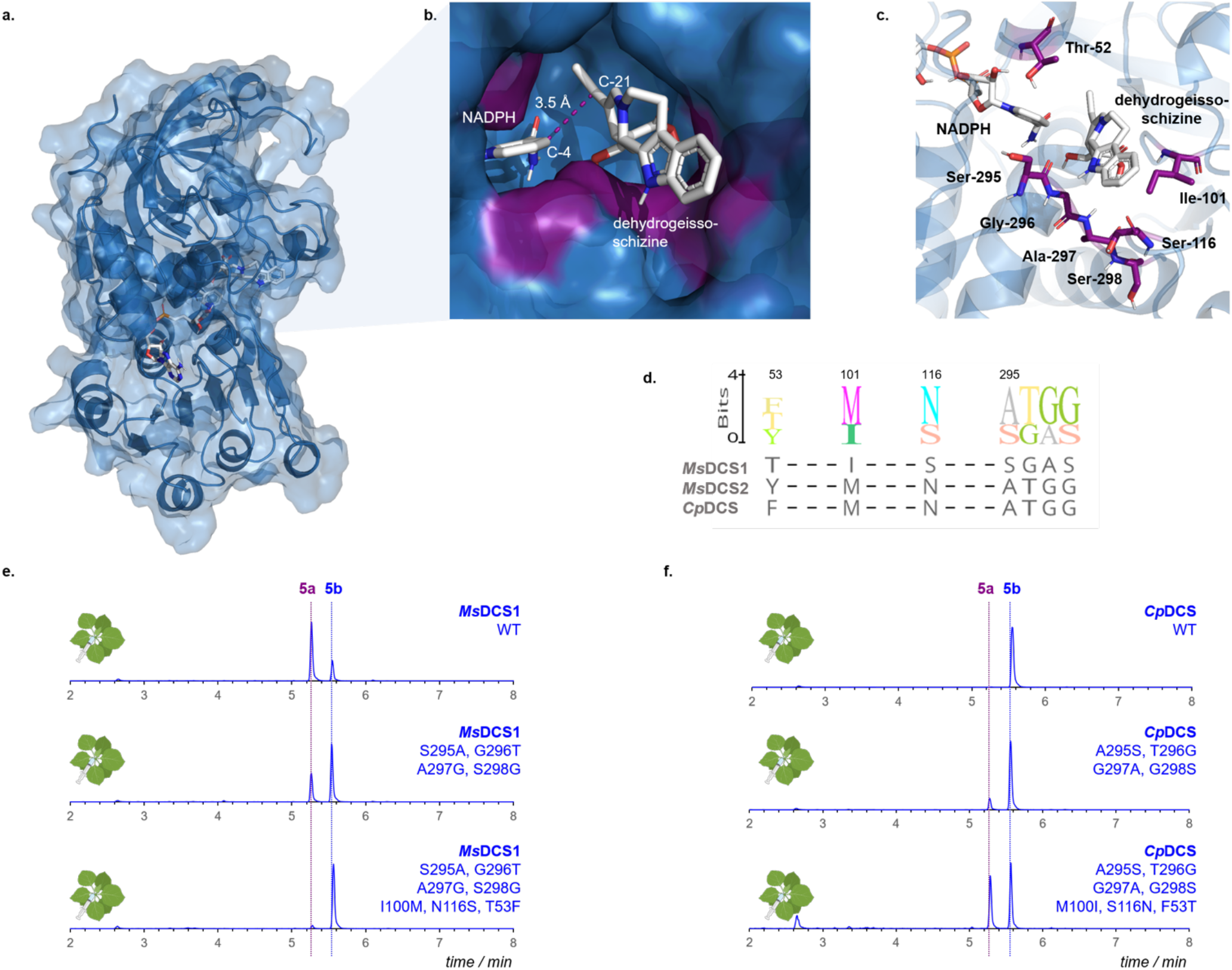
**(a**,**b)** Structural model of *Ms*DCS1 in complex with NADPH and dehydrogeissoschizine (**15**); (**c**) Key active site residues directing the stereoselectivity; **(d)** Sequence alignment between *Ms*DCS1, *Ms*DCS2 and *Cp*DCS; **(e,f)** Levels of (20*S*)-**5a** and (20*R*)-**5b** in mutants of *Ms*DCS1/*Cp*DCS; displayed are EIC (*m/z* 369) corresponding to transient expression of *Cr*STR, *Cr*SGD, ADH-mutants and *Ms*EnolMT in *Nicotiana benthamiana*.

These seven amino acids may collectively affect the orientation in which dehydrogeissoschizine (**15**) binds in the enzyme active site, which would in turn control tautomerization and protonation of **16** to either (20*S*)-**17a** or (20*R*)-**17b** (Scheme 2b). The model of the active site did not contain an amino acid that would be appropriately positioned to catalyze this stereoselective protonation, suggesting that a bound water molecule may be responsible, as previously proposed for other monoterpene indole alkaloid reductases.^21^ This mutational analysis lays the foundation for metabolic engineering strategies to improve production of mitragynine **(1)**; for example, *Ms*DCS2 could be knocked out or mutated in Kratom to generate plants with increased levels of the more pharmacologically important Kratom alkaloids with (20*S*)-stereoconfiguration.

Completion of mitragynine (**1**) and speciogynine (**3**) biosynthesis requires methoxylation at C-9 of **5ab** (Scheme 1b).^22^ Since early pathway genes are expressed in root, while **1** is found exclusively in leaves, it is difficult to predict where the genes responsible for methoxylation would be located. Therefore, we screened oxidases with a variety of tissue expression profiles. However, although 172 candidate oxidase genes were assayed, none showed activity towards either **5a** or **5b** (Supplementary Fig. S13-S14). Attempts to use the fungal cytochrome P450 monoxygenase PsiH, the other known oxidase known to hydroxylate this position of the indole moiety, also failed to hydroxylate **5a** or **5b** (Supplementary Fig. S15).^23^

Therefore, we switched to a mutasynthetic strategy to reconstitute mitragynine (**1**) biosynthesis. *N. benthamiana* leaves were transiently expressed with *Cr*STR, *Cr*SGD, *Ms*DCS1 and *Ms*EnolMT and infiltrated with commercially available 4-methoxy-tryptamine (**18**) and secologanin (**19**). Consistent with the previously observed stereoselectivity of *Ms*DCS1, this afforded a mixture of **1** and **3**, with **1** as the dominant product (Scheme 3a,b). Exchange of *Ms*DCS1 with *Cp*DCS solely afforded speciogynine (**3**). *In vitro* assays yielded similar results (Supplementary Fig. S16).

**Scheme 3.**
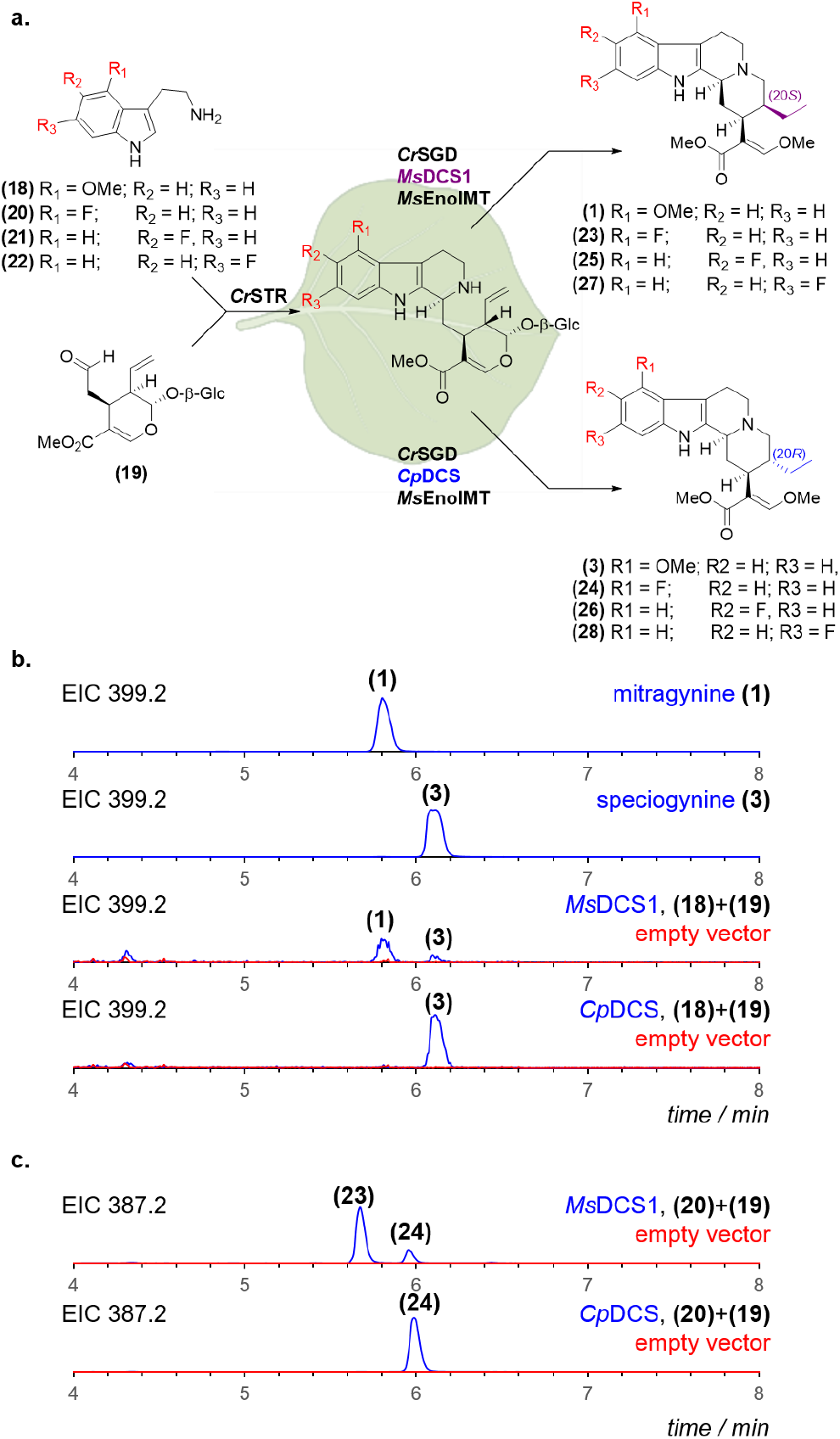
**(a)** *N. benthamiana* infiltration strategy; **(b**,**c)** EIC of methanolic extracts from the transient expression of *Cr*STR, *Cr*SGD, *Ms*EnolMT and indicated ADH enzymes with different tryptamine analogues.

Notably, fluorinated mitragynine analogues have substantially enhanced pharmacological activity.^24^ Therefore, we assessed the potential for the biocatalytic production of fluorinated analogues of **1** and **3**. Infiltration of secologanin (**19**) and either 4F-, 5F- or 6F-tryptamine (**20-22**), along with *Cr*STR, *Cr*SGD, *Ms*DCS1 and *Ms*EnolMT in *N. benthamiana* afforded compounds that corresponded to the expected fluorinated analogues (**23-28**) as evidenced by HRMS (Scheme 3a,c; Supplementary Fig. S17-S19). Although attempts to isolate these compounds in quantities sufficient for NMR analysis failed, this sets the stage for exploring more efficient yeast based strategies for mitragynine analogue engineering.

In conclusion, we elucidated the key enzymatic steps for the production of corynanthe-type alkaloids in Kratom. Mutagenesis experiments suggest a mechanism that is responsible for the control of the stereochemistry at the crucial C-20 position. These discoveries will enable targeted genome editing in Kratom plants to fine-tune alkaloid profiles. Given the recent advent of yeast expression systems for the production of MIAs,^25^ we anticipate that these enzymes will enable development of robust production platforms for mitragynine and speciogynine and related analogues.

## Abbreviations

ADH: alcohol dehydrogenase
*Cr*: *Catharanthus roseus*
*Cp*: *Cin-chona pubescens*
DCS: dihydrocory-nantheine synthase
EIC: ex-tracted ion chromatogram
FPKM: fragments per kilobase of exon per million of mapped fragments
GES: geraniol synthase
G8H: geraniol 8-hydroxylase
GOR: 8-hydroxygeraniol oxidoreductase
GPP: geranylpyrophosphate
ISY: iridoid synthase
hMOR: human μ-opioid receptor
HRMS: high-resolution mass spectrometry
MIA: monoterpene indole alkaloids
*Ms*: *Mitragyna speciosa*
NADPH: nicotinamide adenine dinucleotide phosphate
SAM: *S*-adenosyl-methionine
SGD: strictosidine glucosidase
STR: strictosidine synthase
TIC: total ion chromatogram

## ASSOCIATED CONTENT

### Supporting Information

Supporting Information is available free of charge on the ACS Publications website as a PDF file. Supplemental information includes detailed methods and materials, supplemental tables (Table S1 – Table S4) and supplemental figures (Figure S1 – Figure S20).

### Accession Codes

GenBank accession numbers: xxxxxxxxx (*Ms*DCS1); xxxxxxxxx (*Ms*DCS2);: xxxxxxxxx (*Ms*EnolMT);: xxxxxxxxx (*Cp*DCS).

## AUTHOR INFORMATION

### Present Addresses

**Yindi Jiang** - CAS Key Laboratory of Quantitative Engineering Biology, Shenzhen Institute of Synthetic Biology, Shenzhen Institute of Advanced Technology, Chinese Academy of Sciences, Shenzhen China.

**Thu-Thuy T. Dang** – Department of Chemistry, Irving K. Barber Faculty of Science, University of British Columbia, Kelowna, British Columbia, Canada.

**Francisco León** – Department of Drug Discovery and Biomedical Sciences, College of Pharmacy, University of South Carolina, Columbia, SC 29208, USA.

**Marco Mottinelli** – Laboratory for Neglected Disease Drug Discovery, College of Science, Department of Chemistry and Chemical Biology, Northeastern University, 02115 Boston, USA.

### Notes

The authors declare no competing financial interests.

## ACKNOWLEDGMENT

We gratefully acknowledge Delia Ayled Serna Guerrero, Sarah Heinicke and Maritta Kunert for assistance with mass spectrometry. Jens Wurlitzer is thanked for help with molecular cloning. Eva Rothe and the MPI-CE greenhouse team is thanked for taking care of plants. Carlos E. Rodríguez López is kindly thanked for help with bioinformatics. Maite Colinas, Prashant Sonawane, Chloe Langley and Matilde Florean are kindly thanked for helpful discussion and advice on methodology. This work was supported by grants from the European Research Council (788301) and the Max Planck Society. A portion of this study was supported by UG3DA048353 and R01DA047855 grants from the National Institute on Drug Abuse and the University of Florida Clinical and Translational Science Institute, which is supported in part by the NIH National Center for Advancing Translational Sciences under award number UL1TR001427. The plant art in Figure 1+2 and Scheme 2+3 was created with BioRender.com.

