## Supplementary Information for "Directed Biosynthesis of Mitragynine Stereoisomers"

This file includes:

| | |
|---|---|
| Material and Methods | S2-S8 |
| Supplementary Figures | S9-S40 |
| Supplementary Tables | S41-S47 |
| Supplementary References | S48-S49 |

#### Materials and Methods

##### Plants and Plant Growth

*Mitragyna speciosa* plants were kept in the greenhouse at 25-27 °C during the day and 24-26 °C during the night, following a 12-h light/12-h dark photoperiod. Relative humidity was kept between 70% and 80%. Plants were propagated via cuttings. Tissue samples of *M. speciosa* were collected from mature plants and snap frozen in liquid nitrogen and stored at -80 °C indefinitely. The same samples were used for RNA extraction and metabolomics (*vide infra*).

*Nicotiana benthamiana* plants were grown on a standard soil mix in the greenhouse. Culture conditions were set to 22 °C, 60% relative humidity and followed a 16-h light/8-h dark photoperiod. Tobacco plants were usually grown for at least 3 weeks but no longer than four weeks prior to infiltration with *Agrobacterium tumefaciens* GV3101. Plant watering was performed periodically as needed.

##### Chemicals

All chemicals used in this study were purchased molecular biology grade or higher from commercial vendors (*Sigma Aldrich*, *Thermo Fischer*, etc.) unless denoted different. Kratom alkaloid standards were obtained from the following sources: mitragynine **(1)** from *Biosynth Ltd.*; speciogynine **(3)**, paynantheine **(6)** and speciocilliatine **(7)** were obtained from *Cayman Chemical*. Strictosidine **(8)** was (bio)synthesized in the course of this study (*vide infra*), as recently reported (Caputi *et al.*);[1] 7OH-mitragynine **(2)** and (20*S*)-corynantheidine **(5a)** were kindly gifted to us by Christopher McCurdy.

##### Codon-optimized gene sequences

Codon-optimized gene sequences were obtained from *Twist Biosciences* for *Ms*DCS1, *Ms*DCS2, *Cp*DCS (ADH genes were optimized for *Escherichia coli*; in case of *Cp*DCS we also obtained a non-codon-optimized sequence for expression in *N. benthamiana*) and PsiH (optimized for *Nicotiana benthamiana*). For codon-optimization the manufacturer´s in-house software was used. Native and codon-optimized sequences are listed below (Table S1+S2).

##### Molecular biology kits

All molecular biology kits were used according to the manufacturer´s instructions, unless specified. The RNeasy Mini Kit (*Qiagen*) was used for RNA extraction (*vide infra*). cDNA was subsequently prepared using Superscript™ IV VILO™ master mix (*Thermo Fischer*). For genes or gene fragments destined for downstream applications the Q5® High-Fidelity 2X Master Mix (*New England Biolabs*) was used for amplification. For colony PCR reactions the One*Taq*® Quick-Load® 2X Master Mix with Standard Buffer (*New England Biolabs*) was used. Gene fragments were purified by agarose gel electrophoresis (1% agarose; 120 V, 40 min) and extracted from the gel using a Zymoclean™ Gel DNA Recovery Kit (*Zymo*). All oligonucleotide primers were synthesized by and obtained from *Sigma Aldrich*. Gene cloning was routinely performed using an

In-Fusion kit (Clontech *Takara*). Plasmid DNA was isolated from bacterial cultures using the Wizard® *Plus* SV Minipreps DNA Purification System kit (*Promega*).

##### Metabolomics on *M. speciosa* tissue

Identical plant samples as used for RNA-Seq were subjected for targeted and untargeted metabolomics. Frozen plant material of mature leaves, young leaves, stems, barks and roots were snap frozen in liquid nitrogen and grounded to a fine powder using mortar and pestle. Fresh tissue weight (100 mg) were mixed with 300 µL MeOH (supplemented with 0.1% formic acid) and vortexed vigorously for 1 min. Afterwards the samples were sonicated for 15 min at room temperature. Cell debris was removed by centrifugation (15000 x g; 20 min). The supernatant was filtered through polytetrafluoroethylene (PTFE) syringe filters (0.22 µm), diluted 1:15 with MeOH (supplemented with 0.1% formic acid) and analysed by UPLC/MS (method 1).

##### RNA purification and sequencing

Total RNA of *M. speciosa* (roots, stem, bark, young leaves, mature leaves) was extracted using the RNeasy Mini Kit (*Qiagen*) according to manufacturer´s instructions. The TURBO DNA-*free*™ Kit (*Thermo Fischer*) was used to remove contaminating DNA. Optionally the RNeasy Mini Kit (*Qiagen*) was used again to improve purification. For each tissue triplicates were prepared. The quality of obtained RNA was analysed using an *Implen* NanoPhotometer® N60. All samples satisfied the necessary requirements for total RNA sequencing (≥ 400 ng; A260/280 = 1.8-2.2; A260/230 ≥ 1.8) and were submitted to Novogene (https://en.novogene.com/) for total RNA sequencing using the company´s standard protocols for library preparation and RNA-Seq. ≥ 30 M raw sequencing reads (Illumina, 150 bp paired-end) were acquired per sample.

##### Coexpression analysis

For gene coexpression analysis the transcriptome provided by the sequencing company (Novogene) was used. Additionally, the raw data was assembled in-house using a standard RNA-Seq. bioinformatics pipeline. In brief, raw read quality was assessed using FastQC (https://www.bioinformatics.babraham.ac.uk/projects/fastqc/).[2] Trimmomatic was used to remove adapter sequences from raw sequencing data.[3] Next, Trinity was used to assemble the *M. speciosa* transcriptome.[4] The transcriptome assembly was refined using the CD-Hit-Suite to group transcripts with greater than 90% identity and only the longest transcript was retained.[5] Transdecoder (https://github.com/TransDecoder/TransDecoder) was deployed to identify candidate coding regions within transcript sequences. Functional annotation was then performed by running Blast against the Uniprot and Pfam-database.[6–8] Finally, Salmon was used to quantify the expression of transcripts.[9] Both the commercially obtained and the in-house generated transcriptomes were used to identify candidate genes. Pearson correlation coefficients were calculated using Microsoft Excel using the expression profiles of *Ms*DCS1 or *Ms*EnolMT as `bait´. Additionally, a self-organising map (SOM) was used to group transcripts by spatiotemporal expression pattern, as recently reported.[10] Genes that grouped with *Ms*DCS1 and *Ms*EnolMT and were functionally annotated as oxidases were considered candidate genes for hydroxylase activity towards corynantheidine **(5ab)**.

#### Cloning of gene candidates

Full-length gene sequences of candidates were amplified by PCR from cDNA of *M. speciosa* using the primers indicated in Supplementary Table S3+S4 and further purified using agarose gel electrophoresis. Each fragment was amplified with suitable overhangs at the 5´ and 3´ end to facilitate In-Fusion cloning into suitable plant or bacterial expression plasmids. In case synthetic gene sequences were subcloned into expression vectors, the commercially obtained oligonucleotide was used as template for the PCR reaction.

For transient gene expression in *Nicotiana benthamiana*, genes of interest were subcloned into a modified binary 3Ω1 vector.[11] 3Ω1 was linearized with the restriction enzyme *Bsa*I (*Thermo Fischer*), purified by kit (DNA Clean & Concentrator™-5, *Zymo*) and fused with the gene of interest by In-fusion cloning (*Clontech Takara*, manufacturer´s instructions). The In-Fusion reaction mixture was transformed into chemically competent *E. coli* TOP10 cells (*Thermo Fischer*) and selected on LB agar supplemented with spectinomycin (200 µg/mL). After 1 d positive transformants were identified by colony PCR using universal sequencing primers (#33, #34) for 3Ω1 (Supplementary Table S4). Plasmids of positive transformants were isolated from overnight cultures (37 °C, 225 rpm, LB+spectinomycin) and correct cloning was confirmed by Sanger sequencing (*Azenta Life Sciences*).

For gene expression in *Escherichia coli*, genes of interest were ligated in frame downstream of a $His_6$-coding sequence of pOPINF vectors linearized with *Hind*III and *Kpn*I. Alternatively, a pOPINM vector (encoding an N-terminal MBP-tag) linearized with *Hind*III and *Kpn*I was used. pOPINF and pOPINM were kindly provided by Ray Owens (Addgene plasmid #26042 & #26044).

#### Mutagenesis

Site-directed point mutations were introduced into the *Ms*DCS1 or *Cp*DCS gene by PCR. The mutagenesis strategy is illustrated in Supplementary Fig. S20. In brief, to introduce mutations into the respective ADH gene the gene was amplified in two fragments, containing the mutation(s) in complementary overhangs. Both fragments were purified by agarose gel electrophoresis (1%, 120 V, 45 min) and ligated into linearized 3Ω1 vector by means of In-Fusion cloning (*Clontech Takara*). Correct cloning was assessed by Sanger sequencing and only sequence verified plasmids were used in downstream applications.

#### Transformation of *Agrobacterium tumefaciens* GV3101

Electrocompetent cells of *Agrobacterium tumefaciens* GV3101 (*Goldbio*) were thawed on ice and mixed with plasmid DNA (300-600 ng) that had been checked by Sanger sequencing. After incubation on ice for 30 min, the cell suspension was transferred to an electroporation cuvette and cells were electroporated using a MicroPulser™ (*BioRad*). Cells were mixed with 1 mL LB medium and incubated at 28 °C/225 rpm for 3 h prior to plating on LB agar plates (supplemented with 20 µg/mL rifampicin, 50 µg/mL gentamycin and 200 µg/mL spectinomycin). Plates were kept at 28 °C for 2 d. Single colonies were used to inoculate liquid cultures. Liquid cultures were prepared as 10-20 mL cultures (supplemented with 20 µg/mL rifampicin, 50 µg/mL gentamycin and 200 µg/mL spectinomycin) and cultivated at 28 °C and 250 rpm for up to 24 h. 50 % glycerol stocks were prepared thereof, snap frozen in liquid nitrogen and stored at –80 °C indefinitely.

##### Transient expression of gene candidates in *Nicotiana benthamiana*

Transient expression of gene candidates in *N. benthamiana* was performed as previously reported by Hawes *et al.*[12] In brief, transformed *Agrobacterium* GV3101 strains containing the gene construct of interest were cultivated in 10 mL LB medium (supplemented with 20 µg/mL rifampicin, 50 µg/ml gentamycin and 200 µg/ml spectinomycin) for 16 h at 28 °C and 250 rpm. Afterwards the cells were collected by centrifugation (3000 x g, 30 min) and washed with 1-3 mL infiltration buffer (50 mM MES, 2 mM $Na_3PO_4$, 27.8 mM glucose, 100 µM acetosyringone). After centrifugation (3000 x g, 10 min) the wash step was repeated. Finally, cells were resuspended in 10-15 mL infiltration buffer and the optical density $OD_{600}$ was measured. Upon infiltration of a single *Agrobacterium* strain the suspension was diluted to a final $OD_{600}$ = 0.3 in a total volume of 15 mL. Upon infiltration of multiple *Agrobacterium* strains the strains were diluted so that the final $OD_{600}$ was < 1 (equal concentration for each strain). Resulting suspensions were incubated at RT for 1 h and then infiltrated into the underside of 3-4 week old *N. benthamiana* leaves using a needleless 1 mL syringe. After 3 days the substrate(s) (usually 700 µM tryptamine [or methoxylated/fluorinated tryptamine analogues] and 700 µM secologanin dissolved in 1 mL dd$H_2O$) were infiltrated into the underside of the same leaves previously infiltrated with the *Agrobacterium* strains of choice. At 2 days post-infiltration, leaves were harvested (ca. 100-150 mg fresh weight) and snap-frozen in liquid nitrogen. Each individual infiltration experiment was tested at least 2x times, with biological replicates consisting of at least two leaves from two different tobacco plants.

To assess the different ratios of corynantheidine-formation [**(5a)** *vs* **(5b)**] in different mutants of *Ms*DCS1 and *Cp*DCS*,* each *Agrobacterium tumefaciens* strain harbouring a mutant ADH construct was co-infiltrated with *Cr*STR*, Cr*SGD and *Ms*EnolMT into *N. benthamiana* leaves. For each mutated ADH construct 3x biological replicates were procured, with each replicate consisting of two tobacco leaves infiltrated with *Agrobacteria* and substrate. All wild-type and mutant ADH genes were tested in parallel, using the same batch of *N. benthamiana* to minimize batch effects.

##### Sample harvest and analysis

Harvested, snap-frozen *N. benthamiana* leaf tissue (100 mg) was homogenized on a TissueLyser II (*Qiagen*) using 2 x 2-mm-diameter tungsten beads while shaking vigorously at 22 Hz for 2 min. MeOH (350 µL supplemented with 0.1 % formic acid) was added to each sample, prior to vigorous vortexing for 1 min. After sonication (RT, 15 min) the samples were centrifuged at full speed (>13000 x g, 20 min) and filtered through 0.22 µm PTFE syringe filters. Filtered samples were directly analysed by high-resolution LC-MS and individual metabolites were identified based on comparison of retention times and MS2 spectra with authentic standards. DataAnalysis Version 5.3 (*Bruker*) was used to analyse LCMS data.

<u>Analysis of corynantheidine formation in ADH mutants</u>

To assess the different ratios of corynantheidine-formation [**(5a)** *vs* **(5b)**] in different mutants of *Ms*DCS1 and *Cp*DCS extracted ion chromatograms corresponding to *m/z* = 369 of corynantheidine **(5ab)** were analysed: Peak areas of peaks corresponding to **(5a)** and **(5b)** were determined using the DataAnalysis software and relative percentages of **(5a)** and **(5b)** were calculated. The relative percentages of **(5a)**- and **(5b)**-formation with corresponding standard deviations indicated in Supplementary Figure S11 represent the mean relative percentages calculated from the three biological replicates performed for each mutated ADH construct.

#### Heterologous gene expression in *Escherichia coli*

For production of recombinant enzymes in *E. coli*, expression plasmids containing the gene of interest were transformed into chemically competent *E. coli* BL21(DE3) cells using a standard heat-shock protocol. For the expression of *Catharanthus roseus* strictosidine synthase (*Cr*STR) as well as *Catharanthus roseus* strictosidine glucosidase (*Cr*SGD) previously reported expression constructs were used.[13–15]

Single colonies from these transformations were inoculated into a 10 mL seed culture (LB-medium; supplemented with either 100 µg/mL carbenicillin or 50 µg/mL kanamycin) and cultivated overnight at 37 °C and 200 rpm. An aliquot of the seed culture (8 mL) was used to inoculate 1 L 2TY medium (supplemented with either 100 µg/mL carbenicillin or 50 µg/mL kanamycin). The resulting expression culture was incubated at 37 °C and 200 rpm until an optical density ($OD_{600}$) of between 0.4-0.6 was reached. The culture was then moved to an 18 °C shaker set to 200 rpm and protein expression was induced by adding isopropyl β-D-1-thiogalactopyranoside (IPTG) to a final concentration of 200 µM. The culture was incubated overnight for 16-24 h. The cells were then harvested by centrifugation (4000 rpm, 4 °C, 20 min), frozen in liquid nitrogen and stored indefinitely at –80 °C.

#### Purification of recombinant proteins

Cell pellets were thawed on ice and resuspended in 80-100 mL buffer A (50 mM Tris base, 50 mM glycine, 500 mM NaCl, 20 mM imidazole, 5% glycerol (v/v), pH 8.0; a fresh 100 ml aliquot of this was prepared on the day of protein purification and mixed with 10 mg lysozyme and 1x protease inhibitor cocktail tablet [cOmplete, EDTA-free, *Roche*]). Resuspended cells were lysed using an ultrasonic liquid processor (vibra cell™, *Sonics*®; 40 % amplitude; 2s on/3s off; total \`on´-time: 3 min). Cell debris was removed by centrifugation (4 °C, 35 min, 35000 x g). The protein of interest was then purified on an ÄKTA pure FPLC system (*GE Healthcare*) connected to a HisTrap™ column (*cytiva*, column volume = 5 mL). The FPLC system was programed to: [**A**], equilibrate the column [flow rate = 5 ml/min] with 5x column volumes of buffer A (50 mM Tris base, 50 mM glycine, 500 mM NaCl, 20 mM imidazole, 5% glycerol (v/v), pH 8.0); [**B**] load the protein sample [flow rate = 2 ml/min]; [**C**] wash the column [flow rate = 5 ml/min] with buffer A until the UV absorption at 280 nm is stable (stability time = 1 min; accepted UV fluctuation = 0.1 mAU; maximum wash volume = 20x column volumes); [**D**] elute the protein with 5x column volumes of buffer B (50 mM Tris base, 50 mM glycine, 500 mM NaCl, 500 mM imidazole, 5% glycerol (v/v), pH 8.0). Elution of the protein of interest was monitored using UV absorption at 280 nm. Fractions of interest were assessed by SDS gel electrophoresis, pooled and rebuffered to buffer C (20 mM HEPES, 150 mM NaCl, pH 7.5).

#### Enzymatic *in vitro* assays

Enzymatic assays for reductase activity

Recombinant alcohol dehydrogenase (ADH) candidates were assessed using strictosidine **(8)** as substrate. Reaction mixtures (50 µL total volume, 50 mM HEPES, pH 7.4) comprised 1 µM *Catharanthus roseus* strictosidine glucosidase (*Cr*SGD; to catalyse deglycosylation of strictosidine *in situ*), 1 µM of the respective ADH enzyme, 100 µM NADPH and 40 µM strictosidine **(8)**. Assays were kept at 30 °C/400 rpm for 16 h. Negative controls consisted of boiled enzymes (90 °C, 10 min). For LCMS analysis assays were mixed with equal volumes of MeOH, centrifuged at 15000 x g for 20 min and filtered through 0.22 µm PTFE syringe filters prior to untargeted metabolomics (LCMS Method 1).

Enzymatic assays for *O*-methyltransferase activity

Recombinant *Ms*EnolMT candidates were assessed using strictosidine **(8)** as substrate. Reaction mixtures (50 µL total volume, 50 mM HEPES, pH 7.4) were composed of 1 µM *Catharanthus roseus* strictosidine glucosidase (*Cr*SGD; catalysing deglycosylation of strictosidine *in situ*), 1 µM of either *Ms*DCS1/*Ms*DCS2/*Cp*DCS, 2 µM of the respective *Ms*EnolMT candidate, 100 µM NADPH, 200 µM SAM, 200 µM ascorbate and 40 µM strictosidine **(8)**. For LCMS analysis assays were mixed with equal volumes of MeOH, centrifuged at 15000 x g for 20 min and filtered through 0.22 µm PTFE syringe filters prior to untargeted metabolomics (LCMS Method 1).

#### Preparative scale *in vitro* reactions for product isolation

Preparation of strictosidine **(8)**

Strictosidine **(8)** was produced as reported recently by Caputi *et al.*[1] In brief, 6 mM tryptamine hydrochloride and 4 mM secologanin were combined in a total volume of 15 mL HEPES buffer (50 mM, pH 7.5). *Cr*STR was added to a final concentration of 5 µM and the reaction was stirred at 30 °C for 18 h. Initially, strictosidine was pre-purified on a reverse-phase solid phase extraction (SPE) cartridge (Discovery DSC-18, 1g, *Supelco*). To do so the column was activated with 4 ml of MeOH and equilibrated with 4 mL of water. The sample was then loaded onto the column and 4 mL of water were used to wash the column. Elution of strictosidine was achieved with 8 mL of MeOH. Strictosidine was further purified by preparative HPLC (Method 2; *vide infra*). Strictosidine (2.0 mg) was obtained.

#### LCMS data acquisition

Method 1

All compounds and extracts used in this study were analysed using method 1. Method 1 has been previously reported by Kamileen *et al.*[16] In brief, for LCMS data acquisition an UltiMate 3000 ultra-high performance liquid chromatography system (UHPLC; *Thermo Fischer*) connected to an Impact II UHR-Q-ToF (Ultra-High Resolution Quadrupole-Time-of-Flight) mass spectrometer (*Bruker*) was used. Compound separation was achieved using reverse-phase liquid chromatography on a Phenomenex Kinetex XB-C18 (100 x 2.1 mm, 2.6 µm; 100 Å) column operated at 40 °C. Mobile phases: (**A**) water with 0.1 % formic acid; (**B**) acetonitrile; flow rate = 0.6 ml/min. 2 µL sample was injected in each run; authentic standards were prepared as methanol solutions in concentration ranges between 20-100 µM. Chromatography conditions: 10 % **B** for 1 min; 10 % **B** to 30 % **B** in 6 min; 90 % **B** for 1.5 min; 10 % **B** for 2.5 min. Mass spectrometry conditions: mass spectrometry was performed in positive electrospray ionization mode (capillary voltage = 3500 V; end plate offset = 500 V; nebulizer pressure = 2.5 bar; drying gas: nitrogen at 250 °C and 11 L/min). Mass spectrometry data was recorded at 12 Hz ranging from 80 to 1000 *m/z* using data dependent MS2 and an active exclusion window of 0.2 min. Tandem mass spectrometry settings: fragmentation was triggered on an absolute threshold of 400 and restricted to a total cycle time range of 0.5 s; collision energy was deployed in a stepping option model (20-50 eV). To calibrate MS spectrum recording each run was initiated with the direct source infusion of a sodium formate-isopropanol calibration solution (operated by an external syringe pump at 0.18 ml/min using a 5 mL syringe with an ID of 10.3 mm). The initial 1 min of the chromatographic gradient was directed towards the waste.

#### Compound purification using preparative HPLC

Method 2

All compounds isolated in this study were purified using method 2. To this purpose a preparative HPLC system (*Agilent* 1260 Infinity II) equipped with a *Phenomenex* LC column (Luna® 5 µm C18 (2) 100A, 250 x 30 mm, AXIA™ Packed, Ea) and coupled to a multiple wavelength detector and fraction collector was used. As mobile phases **A** (water + 0.1 % formic acid) and **B** (acetonitrile) were used. The flow rate was set to 30 ml $min^{-1}$ and the gradient was as follows: 10-50 % **B** in 33 min, 50 % **B** for 2 min, 10 % **B** for 5 min. Samples were prepared in MeOH as concentrated solutions (1-5 mg $mL^{-1}$), filtered through a 0.22 µm PTFE syringe filter and injected successively (injection volume: 800 µL). All fractions were assessed by LCMS (Method 1) and fractions containing the desired product were pooled and dried using a Genevac EZ-2 Plus (not HCl compatible) evaporation system.

#### Molecular Docking

For molecular docking of the NADPH cofactor and dehydrogeissoschizine into the active site of *Ms*DCS1 AutoDock Vina on the Webina webserver was used.[17] Default parameters were selected. Protein, cofactor and ligand coordinates were converted into PDBQT format using AutoDock Tools v1.5.7.[18] Docking results were assessed manually and ligand orientations were selected so that the 4-pro-*R*-hydride of the NADPH cofactor was in reasonable proximity to C-21 of dehydrogeissoschizine, the site of initial ligand reduction. Note that the depicted orientation does not necessarily correspond to the lowest possible energy solution. Docking Results were visualized using PyMOL.

#### Supplementary Figures

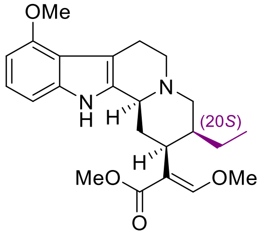

mitragynine **(1)**
Exact Mass: 398.2206

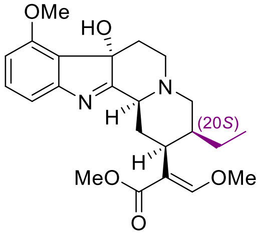

7OH-mitragynine **(2)**
Exact Mass: 414.2155

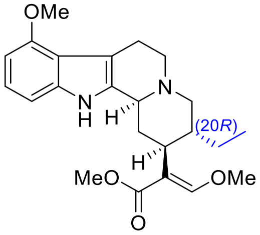

speciogynine **(3)**
Exact Mass: 398.2206

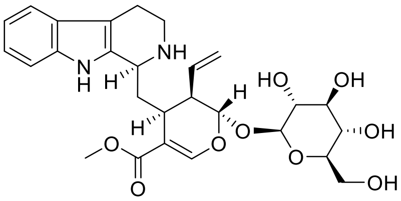

strictosidine **(8)**
Exact Mass: 530.2264

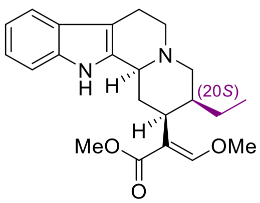

(20*S*)-corynantheidine **(5a)**
Exact Mass: 368.2100

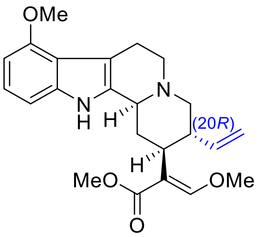

paynantheine **(6)**
Exact Mass: 396.2049

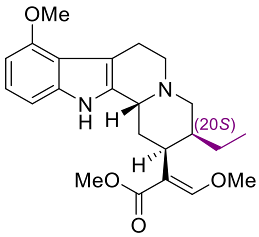

speciocilliatine **(7)**
Exact Mass: 398.2206

**Supplementary Figure S1 | Targeted metabolomics.** Depicted are the chemical structures as well as exact masses of authentic standards used for targeted metabolomics on different kratom tissue material (Supplementary Fig. S2-S6).

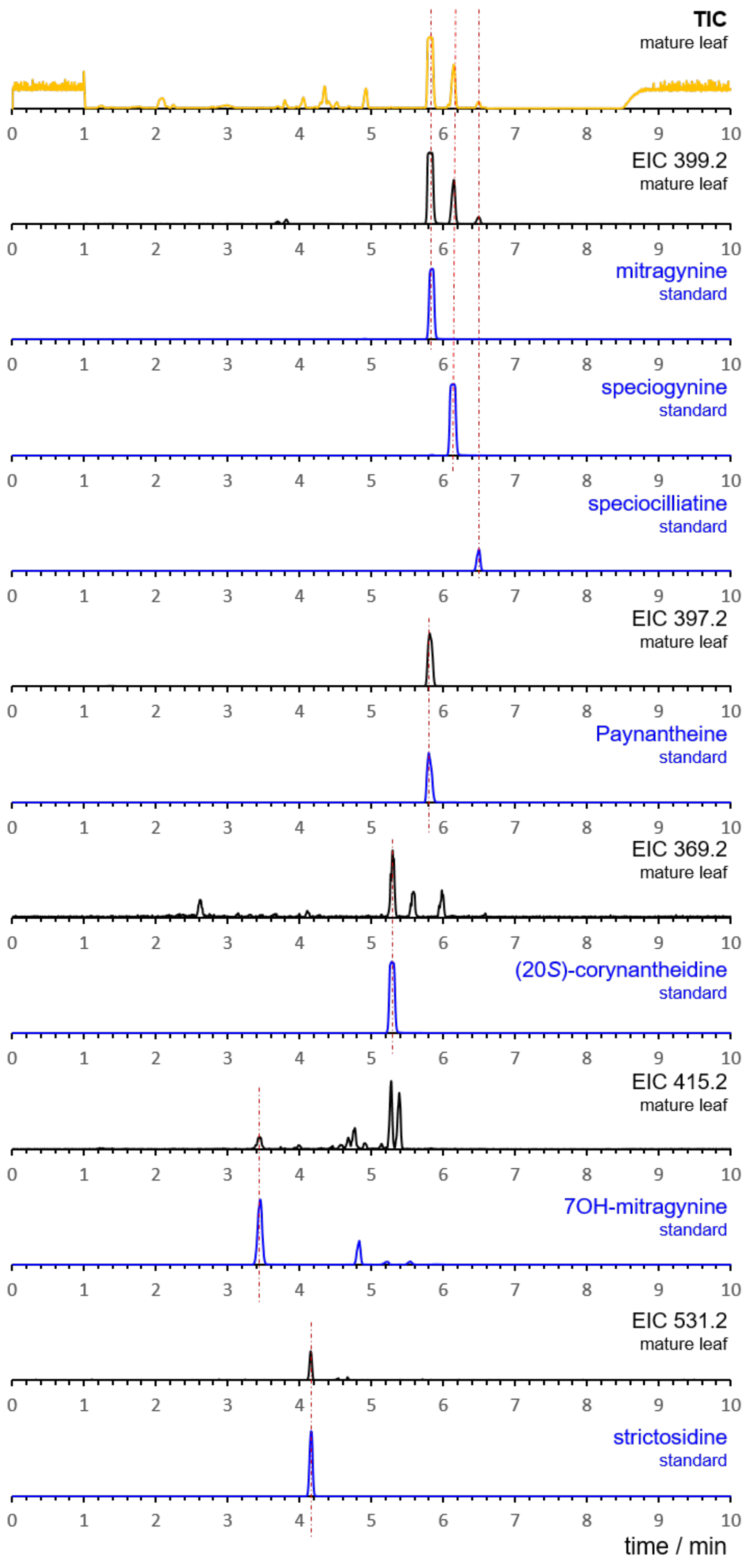


**Supplementary Figure S2 |** See next page for caption.

**Supplementary Figure S2 | Targeted metabolomics on mature leaves of *M. speciosa*.** Methanolic extracts of mature leaves of *M. speciosa* were analysed using LCMS method 1 and compared to authentic standards; top trace (yellow) = total ion chromatogram (TIC) of the mature leaf extract; underneath the TIC trace are shown the extracted ion chromatograms (EIC) of the mature leaf extract that correspond to *m/z* of the standards used in this study; extracted ion chromatograms of authentic standards are shown in blue; metabolites were identified based on identical retention times and identical MSMS data.

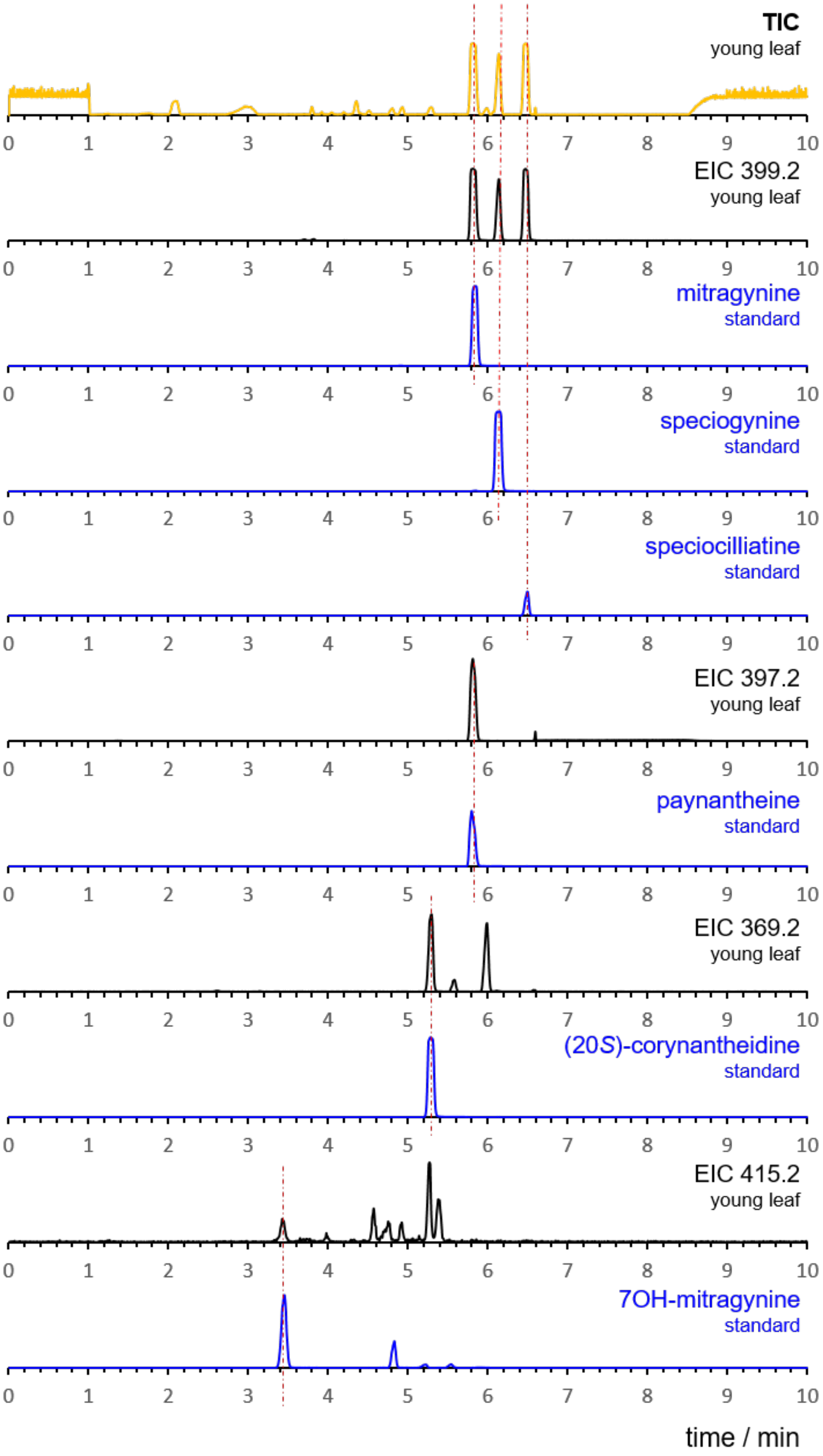


**Supplementary Figure S3** | See next page for caption.

**Supplementary Figure S3 | Targeted metabolomics on young leaves of *M. speciosa*.** Methanolic extracts of young leaves of *M. speciosa* were analysed using LCMS method 1 and compared to authentic standards; top trace (yellow) = total ion chromatogram (TIC) of the young leaf extract; underneath the TIC trace are shown the extracted ion chromatograms (EIC) of the young leaf extract that correspond to *m/z* of the standards used in this study; extracted ion chromatograms of authentic standards are shown in blue; metabolites were identified based on identical retention times and identical MSMS data.

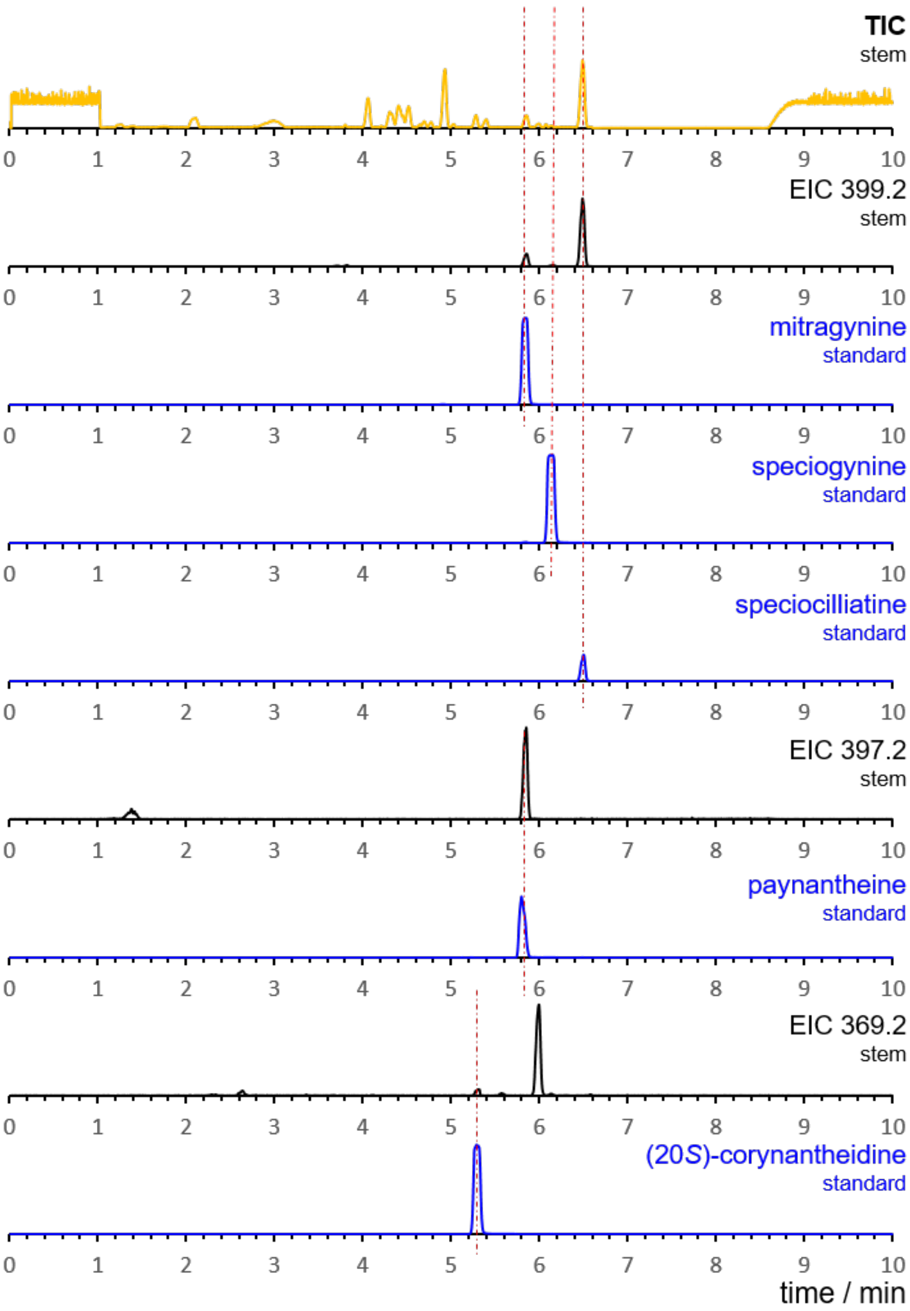


**Supplementary Figure S4 | Targeted metabolomics on stem of *M. speciosa*.** Methanolic extracts of stem of *M. speciosa* were analysed using LCMS method 1 and compared to authentic standards; top trace (yellow) = total ion chromatogram (TIC) of the stem extract; underneath the TIC trace are shown the extracted ion chromatograms (EIC) of the stem extract that correspond to *m/z* of the standards used in this study; extracted ion chromatograms of authentic standards are shown in blue; metabolites were identified based on identical retention times and identical MSMS data.

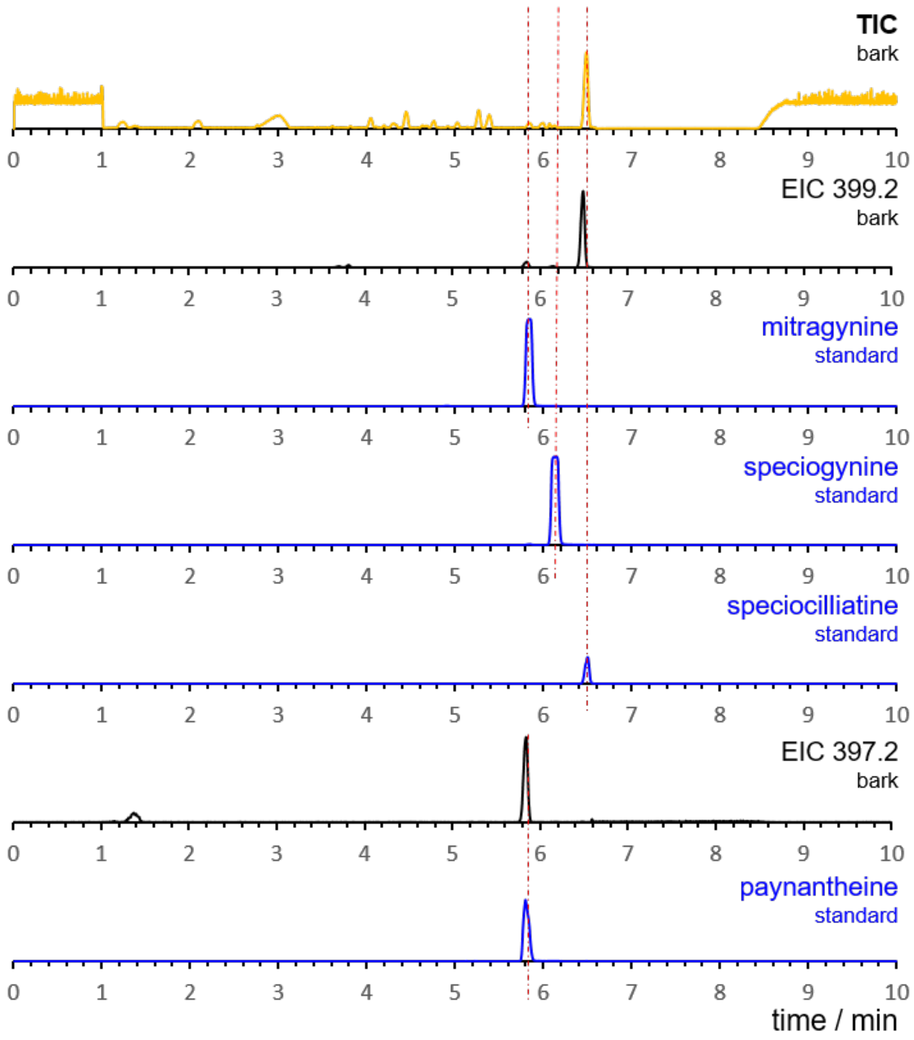


**Supplementary Figure S5 | Targeted metabolomics on bark of *M. speciosa*.** Methanolic extracts of bark of *M. speciosa* were analysed using LCMS method 1 and compared to authentic standards; top trace (yellow) = total ion chromatogram (TIC) of the bark extract; underneath the TIC trace are shown the extracted ion chromatograms (EIC) of the bark extract that correspond to *m/z* of the standards used in this study; extracted ion chromatograms of authentic standards are shown in blue; metabolites were identified based on identical retention times and identical MSMS data.

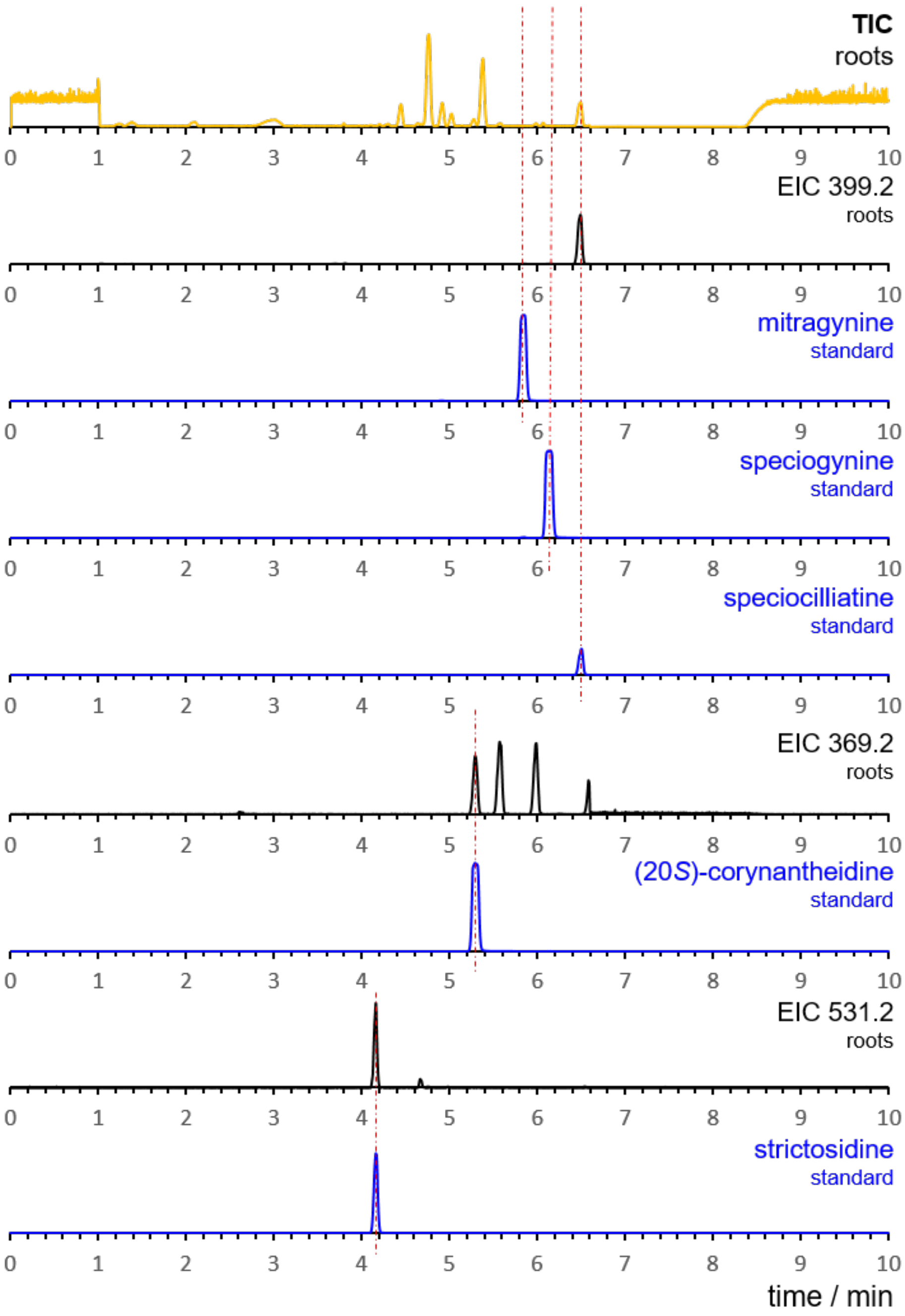


**Supplementary Figure S6 | Targeted metabolomics on roots of *M. speciosa*.** Methanolic extracts of roots of *M. speciosa* were analysed using LCMS method 1 and compared to authentic standards; top trace (yellow) = total ion chromatogram (TIC) of the root extract; underneath the TIC trace are shown the extracted ion chromatograms (EIC) of the root extract that correspond to *m/z* of the standards used in this study; extracted ion chromatograms of authentic standards are shown in blue; metabolites were identified based on identical retention times and identical MSMS data.

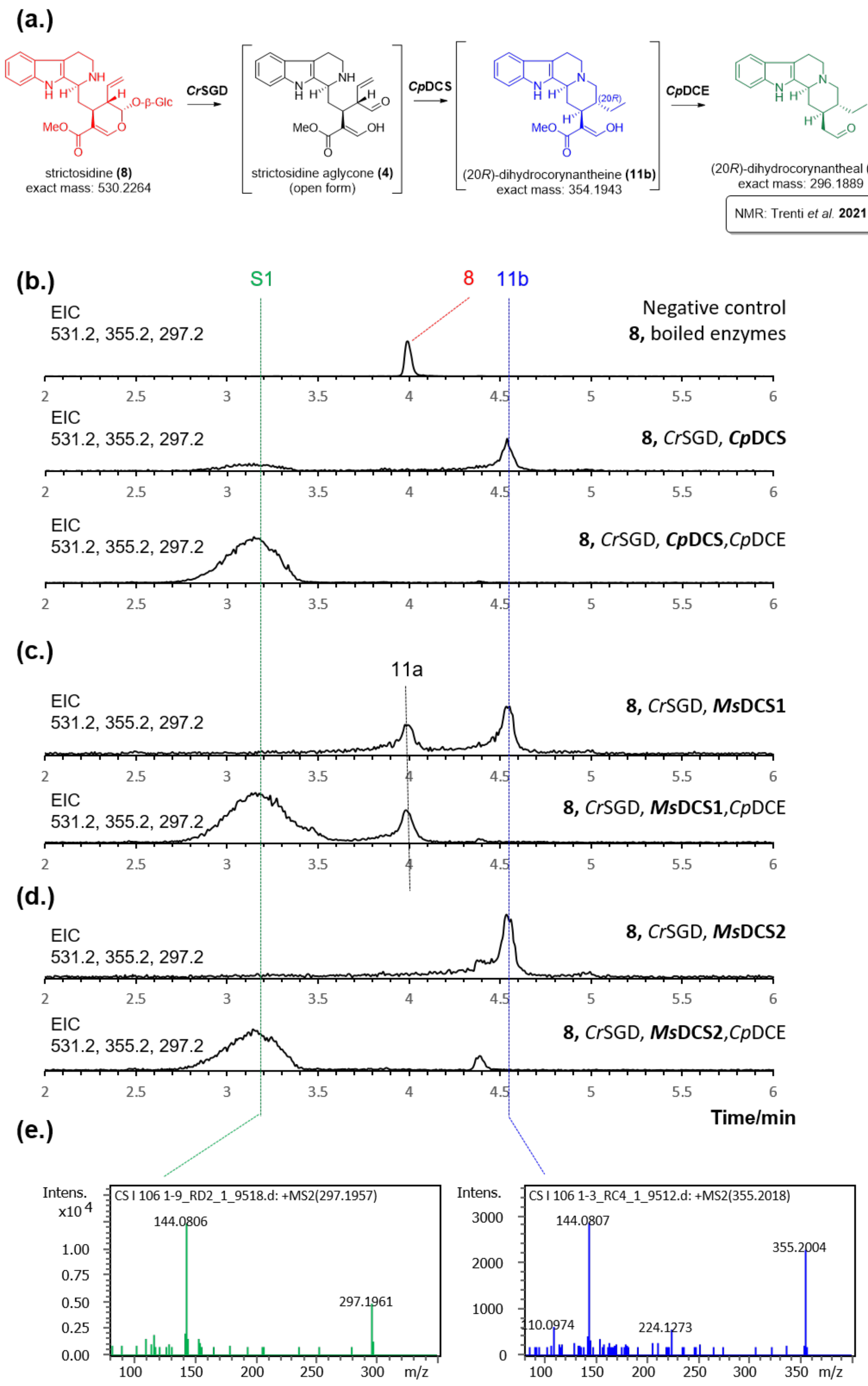


**Supplementary Figure S7** | See next page for caption.

**Supplementary Figure S7 | Characterisation of dihydrocorynantheine synthase products. (a)** Proposed reaction mechanism of dihydrocorynantheine synthase (*Cp*DCS) and dihydrocorynantheine aldehyde esterase (*Cp*DCE).[19] After deglycosylation of strictosidine **(8)** by *Catharanthus roseus* strictosidine glucosidase (*Cr*SGD) the strictosidine aglycone **(4)** gets reduced by *Cp*DCS to (20*R*)-dihydrocorynantheine **(11b)**. Due to the instability of **11b** this product was only identified based on HRMS and MSMS (see panel b,e). Decarboxylation of **11b** is catalyzed by the enzyme dihydrocorynantheine aldehyde esterase (*Cp*DCE) and the stable product **S1** was fully characterized in previous work (Trenti *et. al.*);[19] **(b)** Previous enzymatic *in vitro* assays with *Cp*DCS and *Cp*DCE were repeated in the course of this work; displayed are extracted ion chromatograms (EIC; LCMS method 1) corresponding to the expected *m/z* of strictosidine **(8)** ($[M+H]^+$ = 531.2), dihydrocorynantheine **(11b)** ($[M+H]^+$ = 355.2) and dihydrocorynantheal **(S1)** ($[M+H]^+$ = 297.2); top trace = negative control with **(8)** and boiled enzymes; middle trace = reaction of **(8)** with *Cr*SGD and *Cp*DCS affording a new product corresponding to the formation of **11b**; bottom trace = reaction of **(8)** with *Cr*SGD, *Cp*DCS and *Cp*DCE affords (20*R*)-dihydrocorynantheal **(S1)**, as reported previously; **(c)** Identical *in vitro* assays were performed with *Ms*DCS1 and *Cp*DCE; displayed are extracted ion chromatograms (EIC; LCMS method 1) corresponding to the expected *m/z* of strictosidine **(8)** ($[M+H]^+$ = 531.2), dihydrocorynantheine **(11ab)** ($[M+H]^+$ = 355.2) and dihydrocorynantheal **(S1)** ($[M+H]^+$ = 297.2); top trace = reaction of **(8)** with *Cr*SGD and *Ms*DCS1 affords a mixture of **11a** and **11b**; bottom trace = reaction of **(8)** with *Cr*SGD, *Ms*DCS1 and *Cp*DCE affords (20*R*)-dihydrocorynantheal **(S1); (d)** Identical *in vitro* assays were performed with *Ms*DCS2 and *Cp*DCE; displayed are extracted ion chromatograms (EIC; LCMS method 1) corresponding to the expected *m/z* of strictosidine **(8)** ($[M+H]^+$ = 531.2), dihydrocorynantheine **(11ab)** ($[M+H]^+$ = 355.2) and dihydrocorynantheal **(S1)** ($[M+H]^+$ = 297.2); top trace = reaction of **(8)** with *Cr*SGD and *Ms*DCS2 affording **11b**; bottom trace = reaction of **(8)** with *Cr*SGD, *Ms*DCS2 and *Cp*DCE affords (20*R*)-dihydrocorynantheal **(S1); (e)** HRMS/MS spectra for **11b** and **S1**.

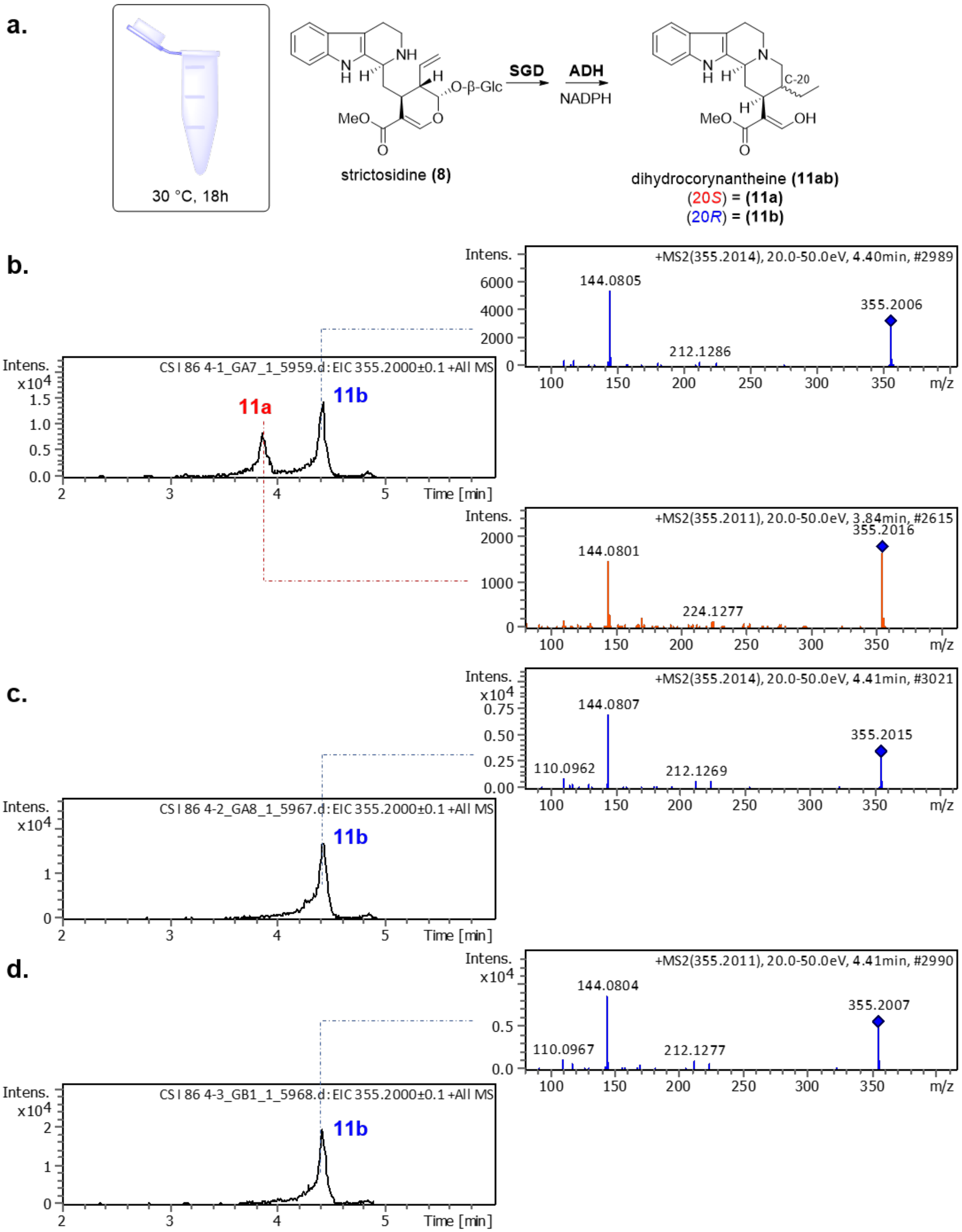


**Supplementary Figure S8** | See next page for caption.

**Supplementary Figure S8 | Identification of (20*S*)-/(20*R*)-dihydrocorynantheine (11ab). (a)** Schematic illustrating enzymatic *in vitro* assays using strictosidine **(8)**, *Catharanthus roseus* strictosidine glucosidase (*Cr*SGD), nicotinamide adenine dinucleotide phosphate (NADPH) and either *Ms*DCS1, *Ms*DCS2 or *Cp*DCS; **(b)** Extracted ion chromatogram and MSMS-data corresponding to *m/z* of **11ab** of assay with *Ms*DCS1; **(c)** Extracted ion chromatogram and MSMS-data corresponding to *m/z* of **11ab** of assay with *Ms*DCS2; **(d)** Extracted ion chromatogram and MSMS-data corresponding to *m/z* of **11ab** of assay with *Cp*DCS.

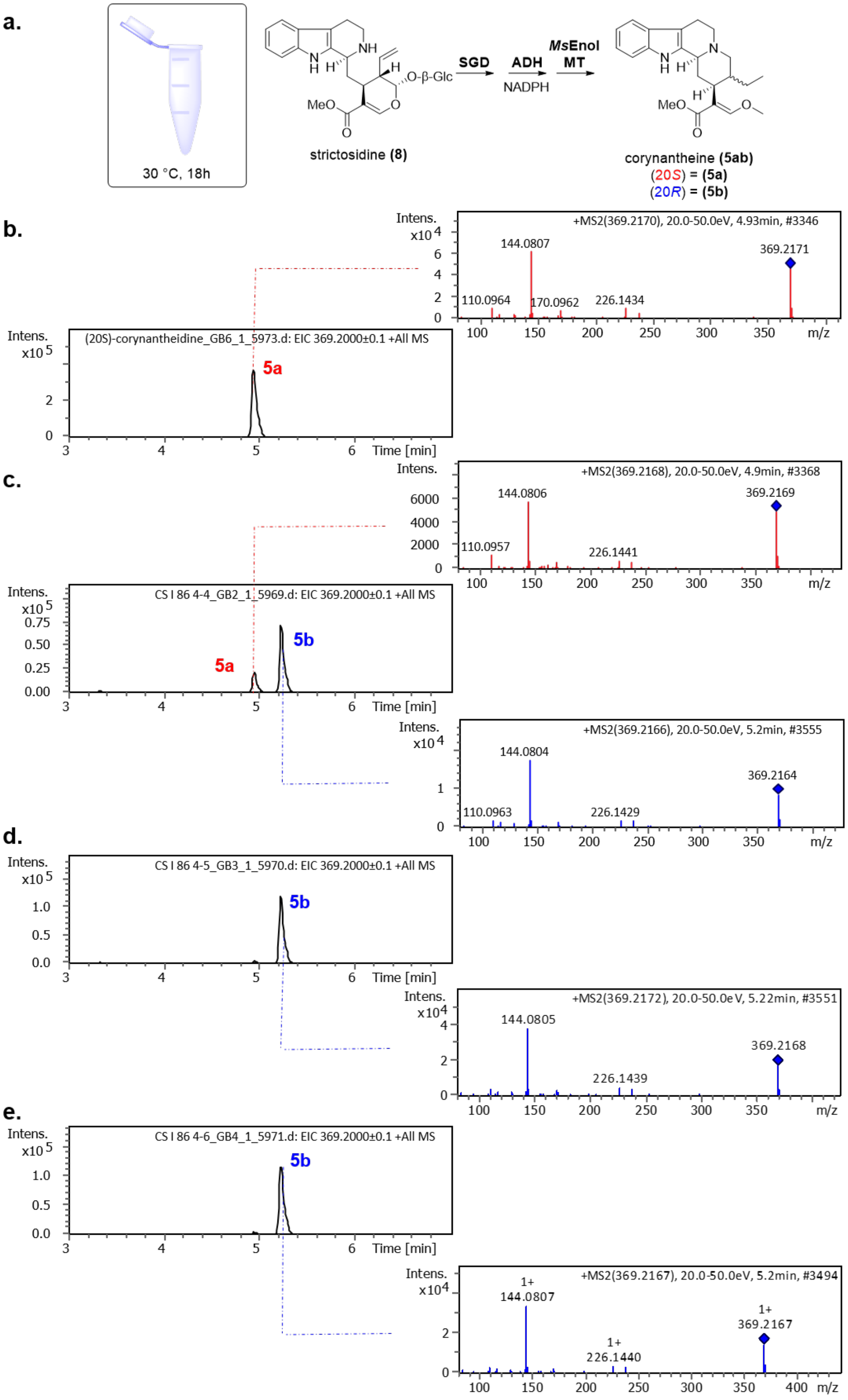


**Supplementary Figure S9** | See next page for caption.

**Supplementary Figure S9 | Identification of (20*S*)-/(20*R*)-corynantheidine (5ab). (a)** Schematic illustrating enzymatic *in vitro* assays using strictosidine **(8)**, *Catharanthus roseus* strictosidine glucosidase (*Cr*SGD), nicotinamide adenine dinucleotide phosphate (NADPH), *S*-adenosylmethionine (SAM), *Ms*EnolMT and either *Ms*DCS1, *Ms*DCS2 or *Cp*DCS; **(b)** Extracted ion chromatogram and MSMS-data corresponding to *m/z* = 369 of standard **5a**; **(c)** Extracted ion chromatogram and MSMS-data corresponding to *m/z* of **5ab** of assay with *Ms*DCS1; **(d)** Extracted ion chromatogram and MSMS-data corresponding to *m/z* of **5ab** of assay with *Ms*DCS2; **(e)** Extracted ion chromatogram and MSMS-data corresponding to *m/z* of **5ab** of assay with *Cp*DCS.

a.

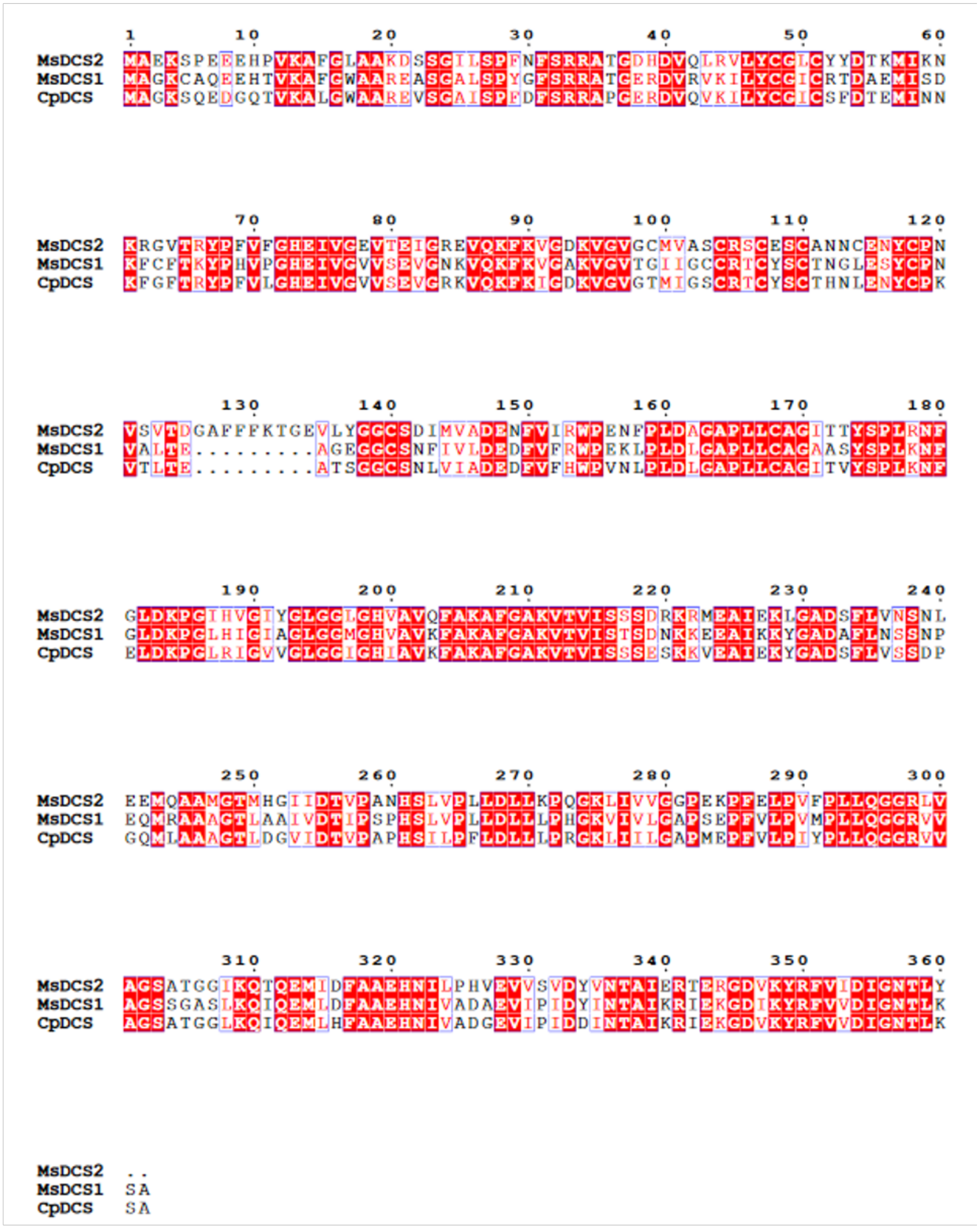


b.

| Enzyme | MsDCS1 | MsDCS2 | CpDCS |
|---|---|---|---|
| MsDCS1 | | 63.4 % | 76.0 % |
| MsDCS2 | 63.4 % | | 64.8 % |
| CpDCS | 76.0 % | 64.8 % | |

**Supplementary Figure S10 |** See next page for caption.

**Supplementary Figure S10 | Amino acid alignment of alcohol dehydrogenases (ADH) used in this study.** **(a)** Protein sequence alignment of *Ms*DCS1, *Ms*DCS2 and *Cp*DCS was created with Clustal Omega;[20] alignment was visualized using ESPript V3;[21] **(b)** Amino acid sequence identity matrix of ADH enzymes used in this study; Muscle 3.8.425 was used for the calculation of sequence identities.[22]

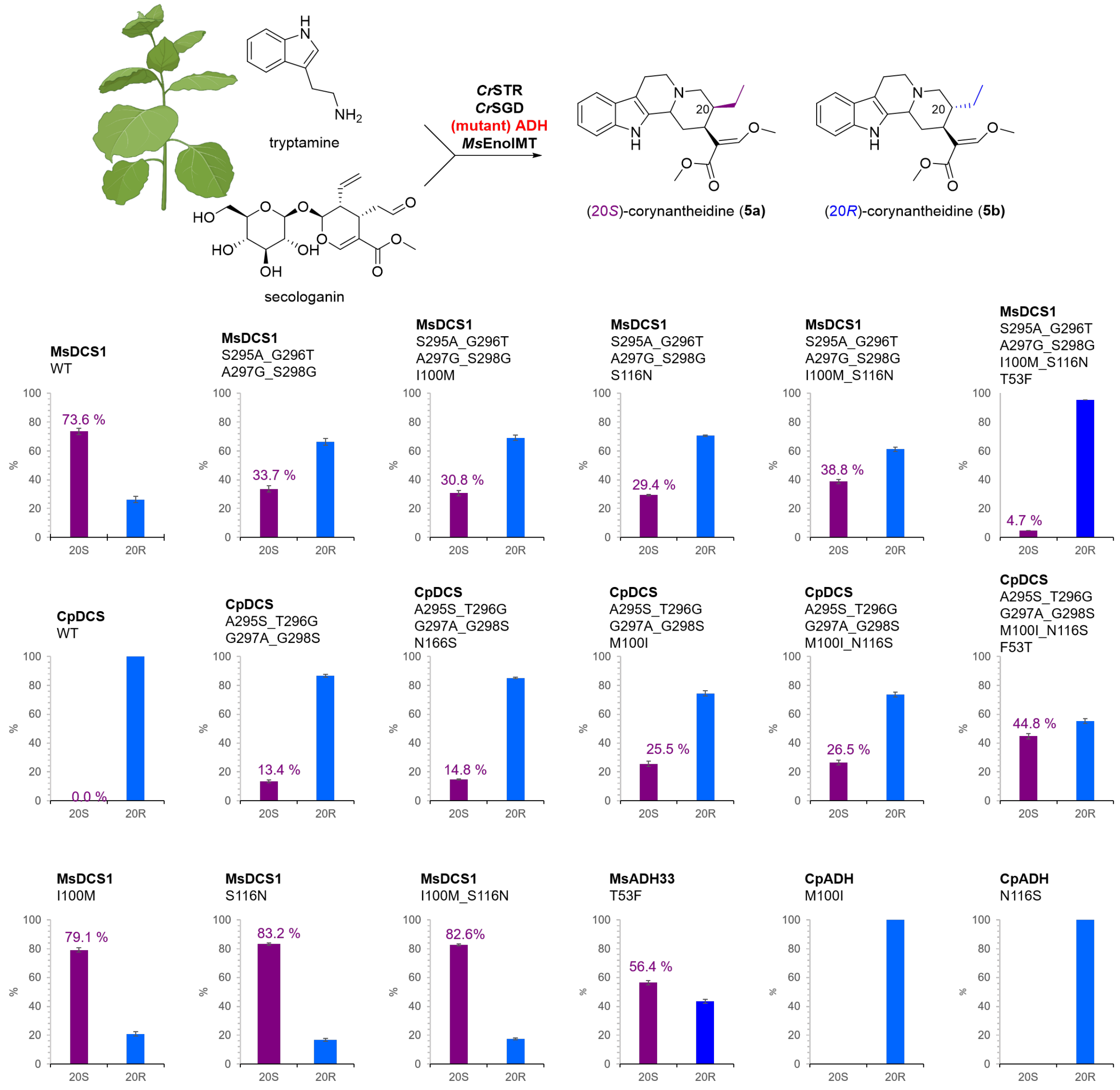


**Supplementary Figure S11 | Effect of key mutants of *Ms*DCS1 and *Cp*DCS on C-20 stereochemistry.** A total of 16 mutants of either *Ms*DCS1 or *Cp*DCS were transiently expressed in *Nicotiana benthamiana*, together with *Catharanthus roseus* strictosidine synthase (*Cr*STR), *Catharanthus roseus* strictosidine glucosidase (*Cr*SGD), *Ms*EnolMT as well as tryptamine (700 µM) and secologanin (700 µM). For each mutant construct, 3x biological replicates were performed. Methanolic extracts were analysed by LCMS (method 1) for the production of (20*S*)-corynantheidine **(5a)** or (20*R*)-corynantheidine **(5b)** and peak areas were determined to calculate the relative percentage of **5a**/**5b**-production. Displayed are the mean relative percentages with corresponding standard deviations calculated from the three biological replicated for each mutated ADH construct.

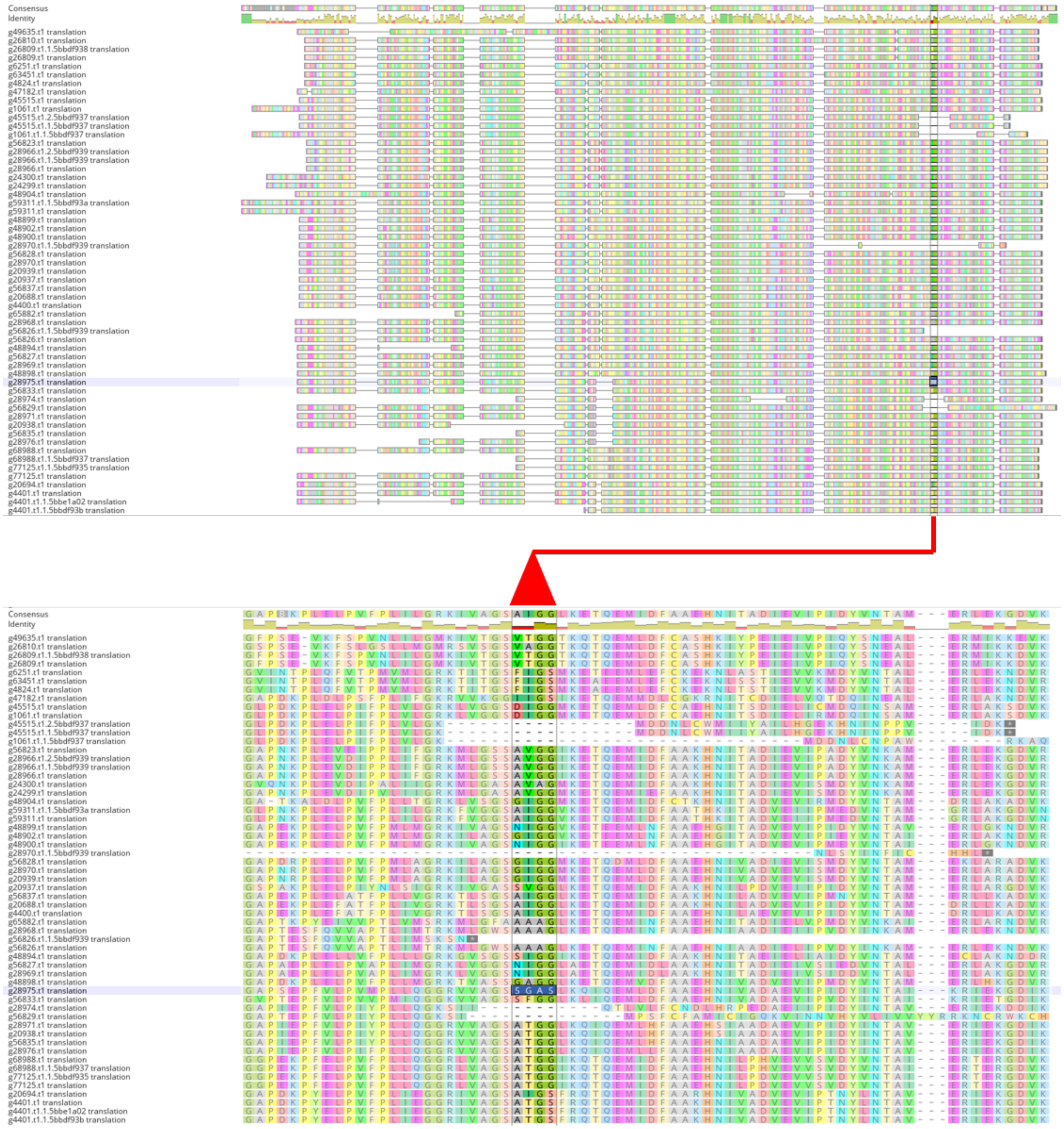


**Supplementary Figure S12 | Mining the kratom genome for *Ms*DCS1 homologues.** Geneious Version 2022.1.1 was deployed to run a local blast of the amino acid sequence of *Ms*DCS1 against the genome of *M. speciosa* (genome has been released by Brose *et al.*;[23] genome downloaded from https://doi.org/10.25387/g3.13042784); ADH homologues were selected based on sequence homology (>40%) and query coverage (>40%) to *Ms*DCS1, affording 57x ADH sequences; Geneious Version 2022.1.1 was used to make a sequence alignment of all sequences (Muscle v3.8.425);[22] a single enzyme (g28975.t1 = *Ms*DCS1; highlighted above) contained the SGAS motif at amino acid position 295-298, which was shown crucial for the formation of the (20*S*)-series of kratom alkaloids.

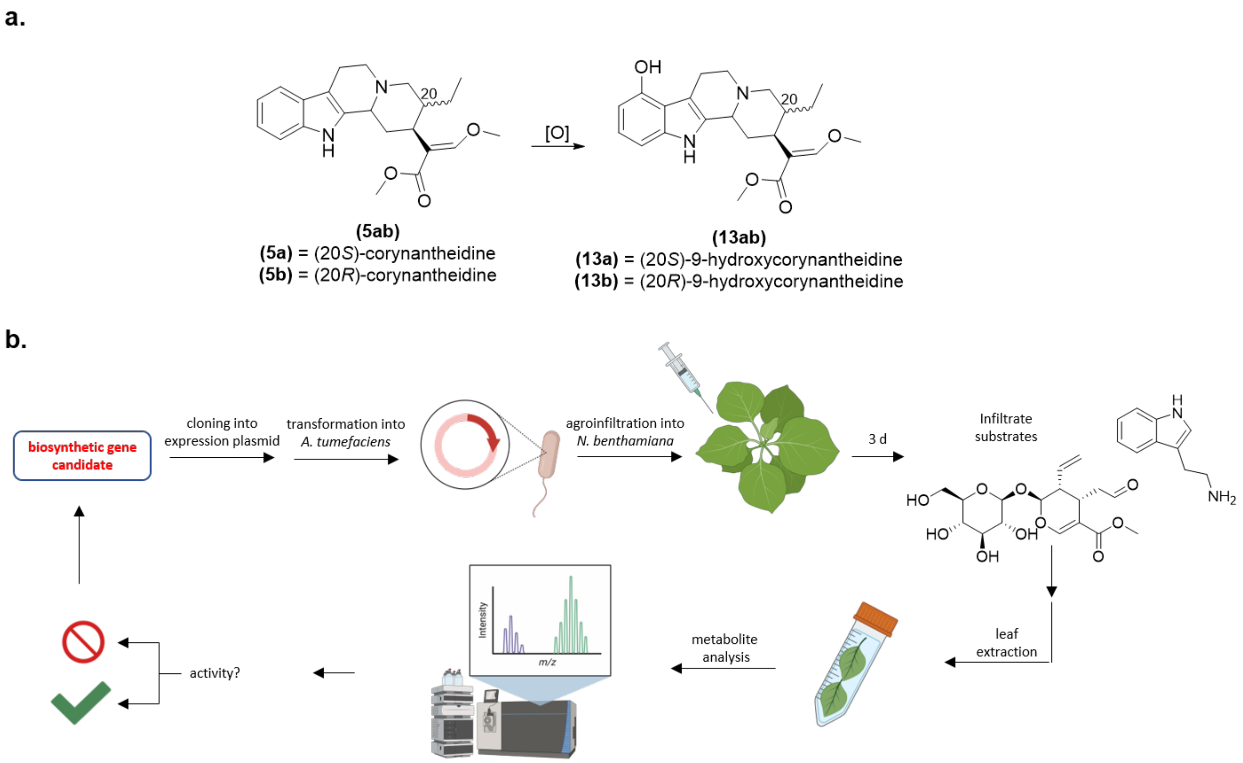


**Supplementary Figure S13 | 9-Hydroxylase screening strategy. (a)** Biosynthetic gene candidates were identified that could catalyze formation of 9-hydroxycorynantheidine **(13ab)** from corynantheidine **(5ab)**; for details of candidate identification see Supplementary Figure S14; **(b)** Biosynthetic gene candidates were subcloned into 3Ω1 vector and transformed into *A. tumefaciens* GV3101. Each candidate was infiltrated together with *Catharanthus roseus* strictosidine synthase (*Cr*STR), *Catharanthus roseus* strictosidine glucosidase (*Cr*SGD), *Ms*DCS1 and *Ms*EnolMT into leaves of *Nicotiana benthamiana*. After 3 days the same leaves were infiltrated with tryptamine (700 µM) and secologanin (700 µM). Due to the transient expression of *Cr*STR, *Cr*SGD, *Ms*DCS1 and *Ms*EnolMT each leaf harbors the potential to produce both epimers of corynantheidine **(5ab)**, the expected biosynthetic precursors for 9-hydroxylation in mitragynine **(1)** and speciogynine **(3)**. After an additional 2 days the infiltrated leaves were harvested, extracted with MeOH and subjected to targeted and untargeted metabolomics.

**a.**

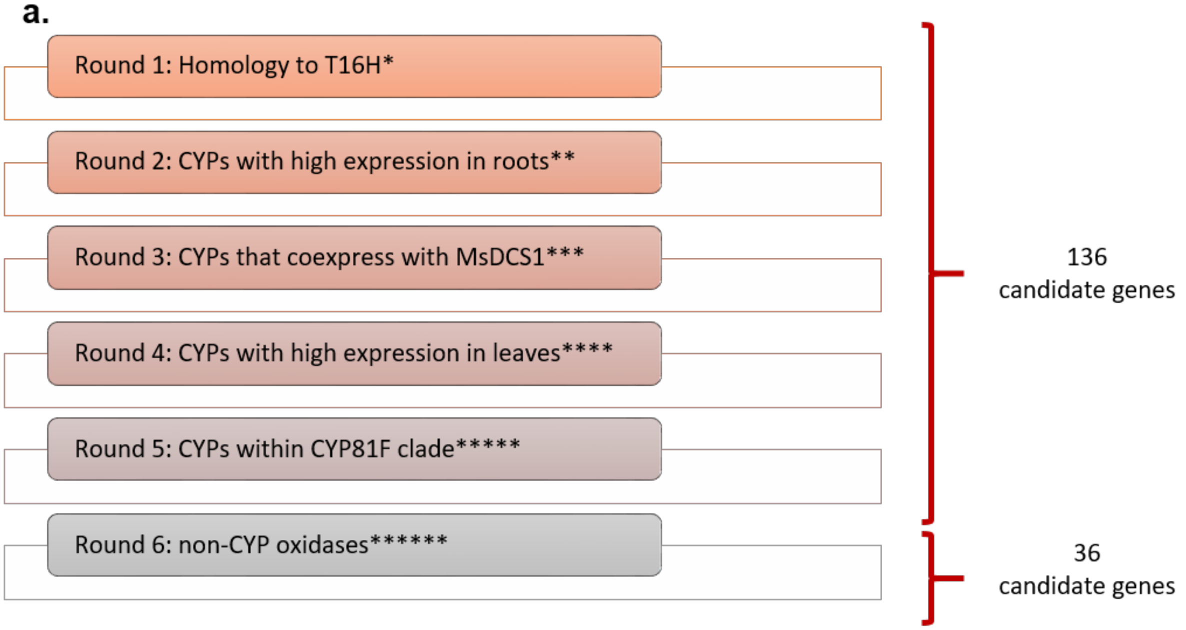


* **<u>Round 1:</u>**…homology to tabersonine-16-hydroxylase (T16H; Uniprot-ID: P98183**);** T16H has previously been shown to hydroxylate the indol moiety of tabersonine (see panel b.)

** **<u>Round 2:</u>**…high expression in the roots of *M speciosa;* iridoid biosynthetic genes and *MsDCS1* / *MsEnolMT* were found to be preferentially expressed in the roots

*** **<u>Round 3:</u>**…coexpression with *MsDCS1* ($r > 0.8$; *Pearson* correlation coefficient)

**** **<u>Round 4:</u>**…high expression in the leaves of *M. speciosa*; metabolomics on *M. speciosa* tissue had revealed that the pathway product mitragynine **(1)** is predominantly found in the leaves

***** **<u>Round 5:</u>**…cytochrome P450 monooxygenases belonging to the CYP81-clade; these have previously been implicated in 4-hydroxylation of the indol-moiety during glucosinolate biosynthesis in *A. thaliana* (see panel c.)

****** **<u>Round 6:</u>**…criteria from round 2-4 were used to identify 7x berberine bridge enzymes, 8x polyphenol oxidases, 2x multicopper-dependent oxidases, 17x 2-oxoglutarate dependent dioxygenases and 2x other oxidases.

**b.**

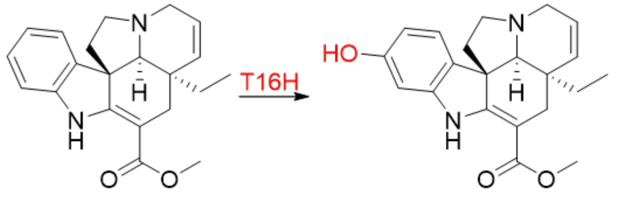


tabersonine 16-hydroxytabersonine

**c.**

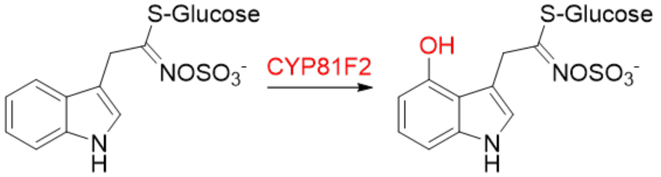


**Supplementary Figure S14 | See next page for caption.**

**Figure S14 | Identification of oxidase genes potentially involved in 9OH-corynantheidine formation. (a)** Selection criteria deployed in six successive candidate identification rounds; **(b)** Hydroxylation reaction catalyzed by T16H, used as bait to identify homologues in kratom;[24] **(c)** Indole hydroxylation in glucosinolate biosynthesis in *Arabidopsis thaliana*.[25]

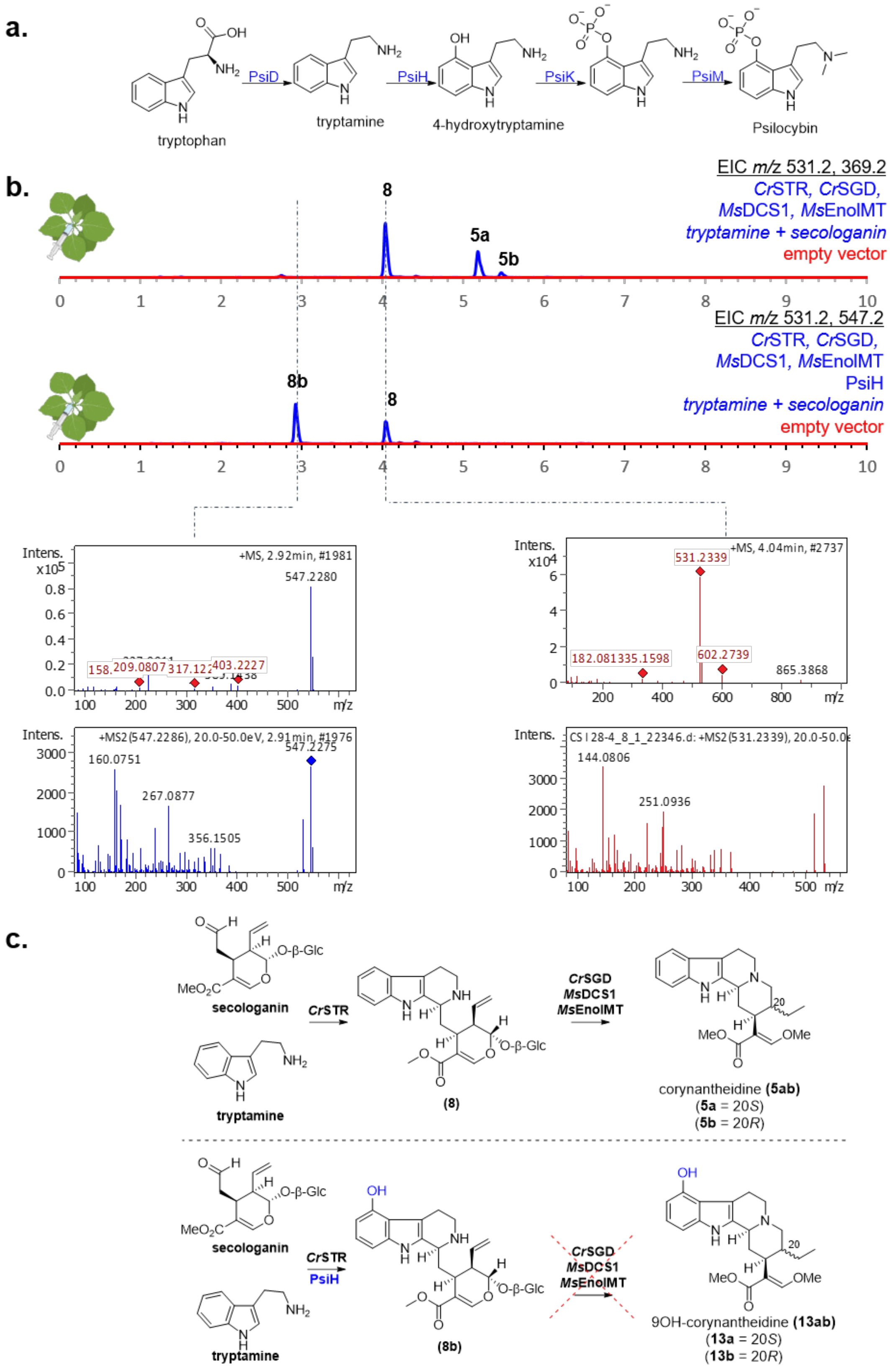


**Supplementary Figure S15 | See next page for caption.**

**Supplementary Figure S15 | Engineering of mitragynine biosynthesis using fungal cytochrome P450 monooxygenase PsiH. (a)** Biosynthetic pathway of psilocybin in *Psilocybe cubensis*;[26] **(b)** Extracted ion chromatograms corresponding to the expected *m/z* of strictosidine **(8)**, 9-OH-strictosidine **(8b)** and corynantheidine **(5ab)**; top trace: transient expression of *Catharanthus roseus* strictosidine synthase (*Cr*STR), *Catharanthus roseus* strictosidine glucosidase (*Cr*SGD), *Ms*DCS1 and *Ms*EnolMT in *Nicotiana benthamiana*; infiltration with tryptamine and secologanin affords strictosidine **(8)** and both epimers of corynantheidine **(5ab)** as identified by HRMS and MSMS data; bottom trace: inclusion of the fungal cytochrome P450 monooxygenase PsiH in the transient expression system leads to formation of a new compound with *m/z* = 547.2 corresponding to the expected formation of 9-hydroxystrictosidine **(8b)**; HRMS and MSMS data are likewise consistent with the formation of **(8b)**; no other new compounds were observed upon transient expression of PsiH, suggesting that one of the downstream enzymes (*Ms*DCS1 or *Ms*EnolMT) do not accept **(8b)** as substrate, implying that hydroxylation occurs after formation of corynantheidine **(5ab)**; **(c)** Top: biosynthetic pathway towards corynantheidine **(5ab)** upon transient expression of *Cr*STR*, Cr*SGD*, Ms*DCS1 and *Ms*EnolMT; bottom: biosynthetic pathway towards 9-hydroxycorynantheidine **(13ab)** in case of early hydroxylation by PsiH; **(13ab)** were not observed in this study, suggesting oxidation occurs after formation of corynantheidine **(5ab)**.

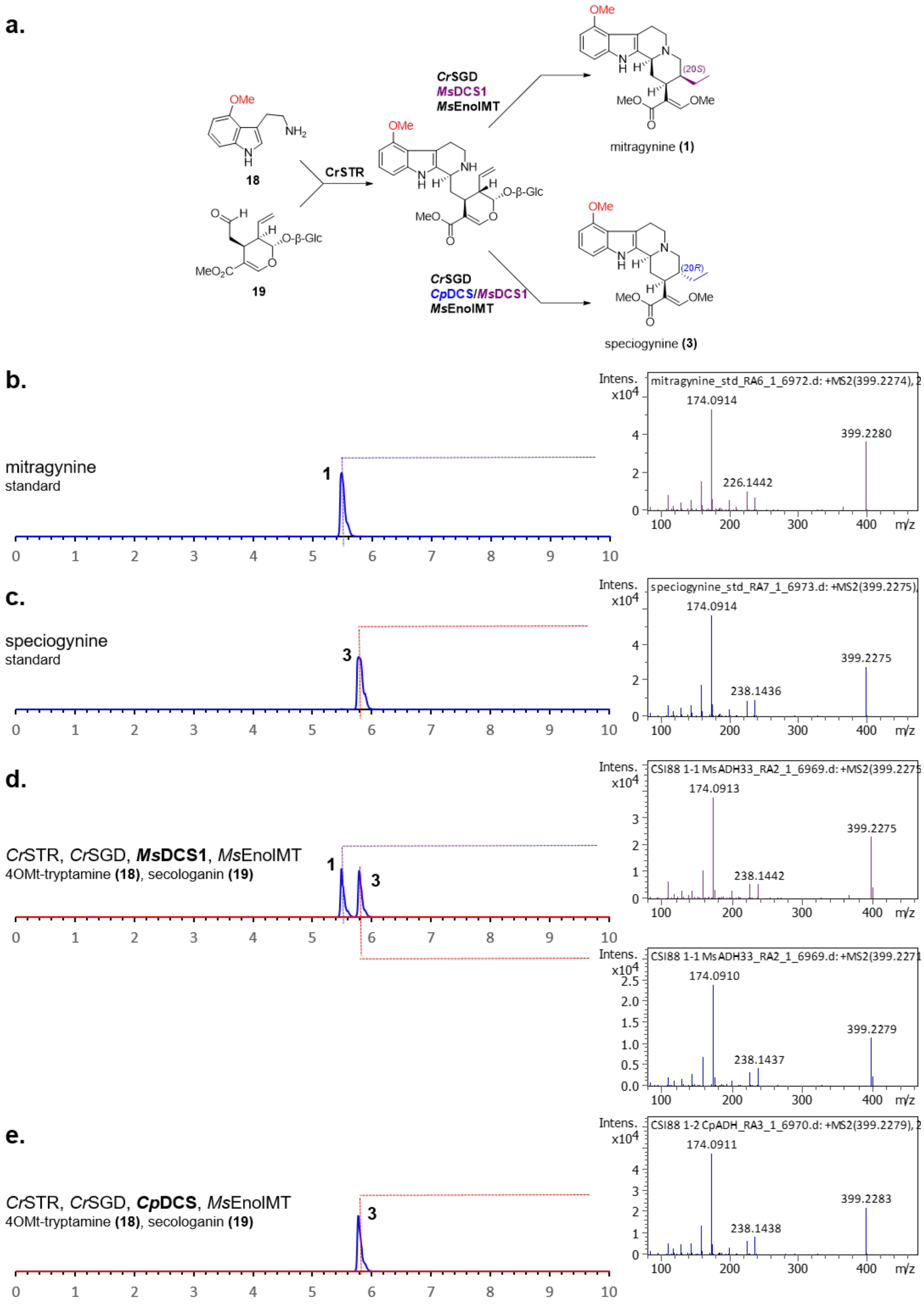


**Supplementary Figure S16 | See next page for caption.**

**Supplementary Figure S16 | Production of mitragynine and speciogynine using recombinant enzymes. (a)** Chosen recombinant enzyme strategy to convert secologanin **(19)** and 4-methoxy-tryptamine **(18)** into mitragynine **(1)** and speciogynine **(3)** using *Catharanthus roseus* strictosidine synthase (*Cr*STR), *Catharanthus roseus* strictosidine glucosidase (*Cr*SGD), *Ms*EnolMT and *Ms*DCS1 or *Cp*DCS; **(b)** Extracted ion chromatogram (EIC; *m/z* = 399) of mitragynine standard and MSMS data; **(c)** Extracted ion chromatogram (EIC; *m/z* = 399) of speciogynine standard and MSMS data; **(d)** Extracted ion chromatogram (EIC; *m/z* = 399) of *in vitro* reaction with recombinant *Cr*STR, *Cr*SGD, *Ms*DCS1, *Ms*EnolMT and the substrates **(18)** and **(19)**; **(e)** Extracted ion chromatogram (EIC; *m/z* = 399) of *in vitro* reaction with recombinant *Cr*STR, *Cr*SGD, *Cp*DCS, *Ms*EnolMT and the substrates **(18)** and **(19)**.

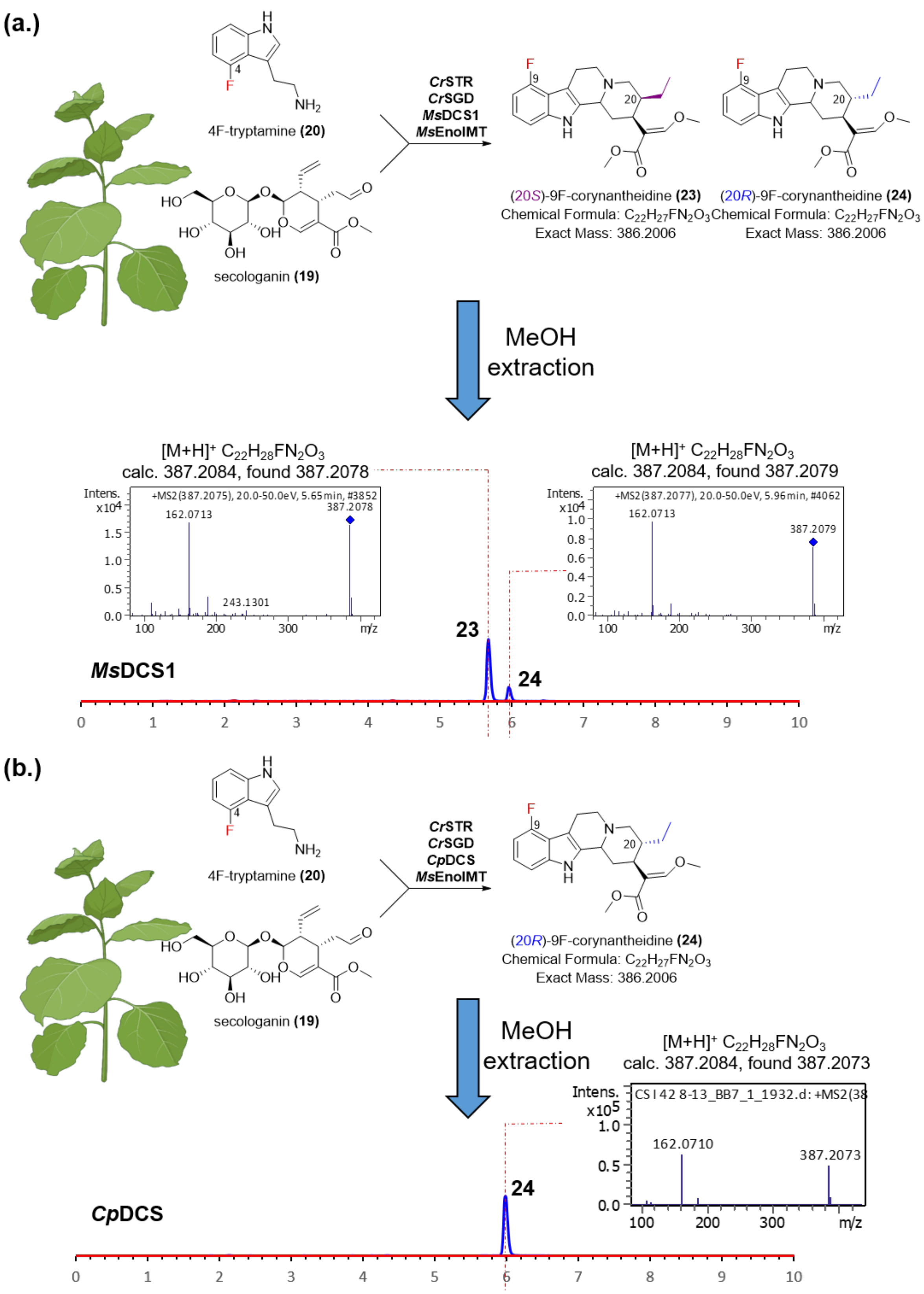


**Supplementary Figure S17 |** See next page for caption.

**Supplementary Figure S17 |** Production of compound corresponding to 9-fluorocorynantheidine. **(a)** *Catharanthus roseus* strictosidine synthase (*Cr*STR), *Catharanthus roseus* strictosidine glucosidase (*Cr*SGD), *Ms*DCS1 and *Ms*EnolMT were transiently expressed in *Nicotiana benthamiana* and infiltrated with 4F-tryptamine **(20)** and secologanin (**19**); methanol extracts were analysed using LCMS method 1; depicted is the extracted ion chromatogram (*m/z* = 387.2) as well as high resolution mass spectrometry and MSMS data corresponding to the formation of (20*S*)-9F-fluorocorynantheidine **(23)** and (20*R*)-9F-fluorocorynantheidine **(24)**; **(b)** *Catharanthus roseus* strictosidine synthase (*Cr*STR), *Catharanthus roseus* strictosidine glucosidase (*Cr*SGD), *Cp*DCS and *Ms*EnolMT were transiently expressed in *Nicotiana benthamiana* and infiltrated with 4F-tryptamine **(20)** and secologanin (**19**); methanol extracts were analysed using LCMS method 1; depicted is the extracted ion chromatogram (*m/z* = 387.2) as well as high resolution mass spectrometry and MSMS data corresponding to the formation of (20*R*)-9F-fluorocorynantheidine **(24)**.

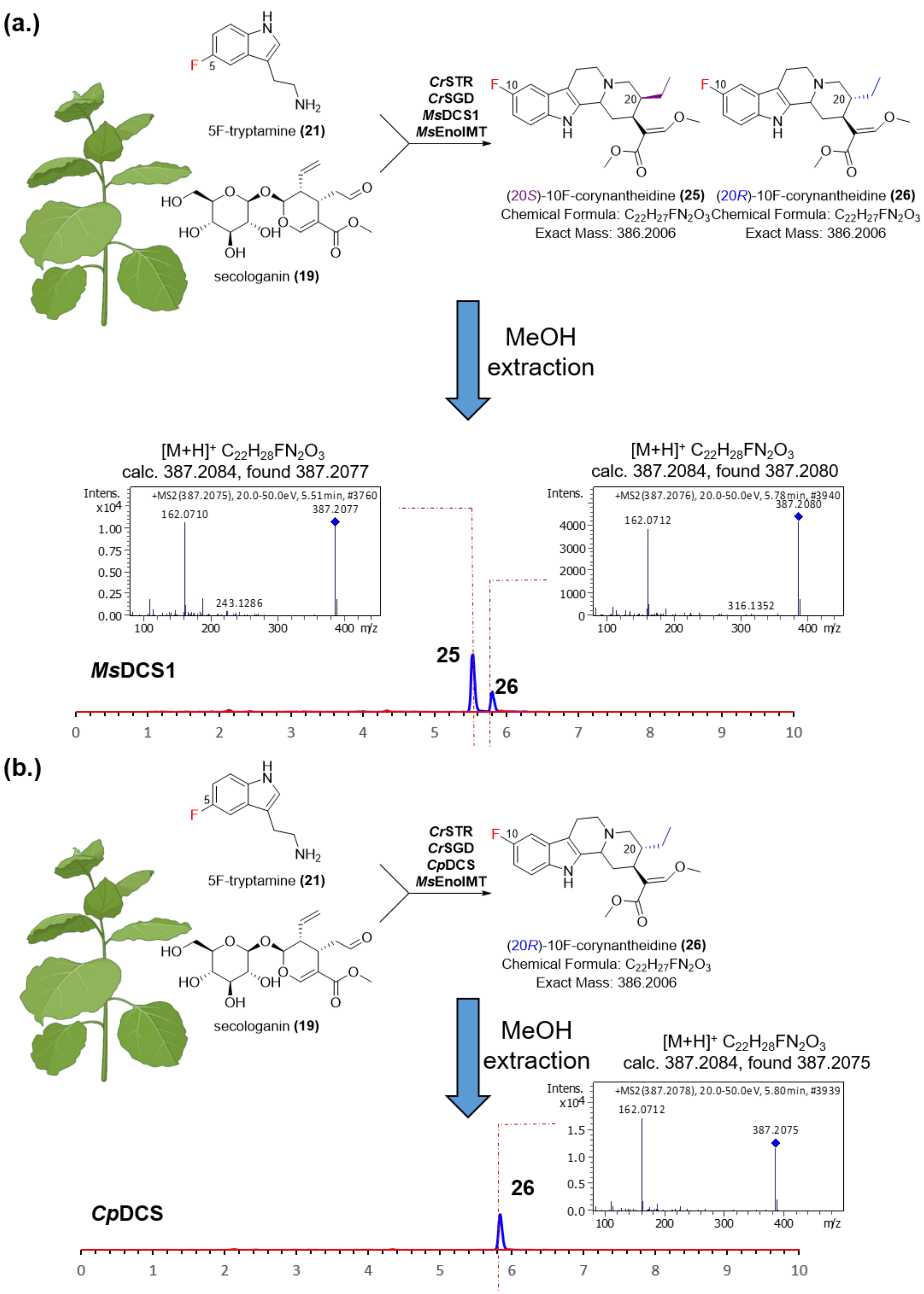


**Supplementary Figure S18 |** See next page for caption.

**Supplementary Figure S18 |** Production of compound corresponding to 10-fluorocorynantheidine. **(a)** *Catharanthus roseus* strictosidine synthase (*Cr*STR), *Catharanthus roseus* strictosidine glucosidase (*Cr*SGD), *Ms*DCS1 and *Ms*EnolMT were transiently expressed in *Nicotiana benthamiana* and infiltrated with 5F-tryptamine **(21)** and secologanin (**19**); methanol extracts were analysed using LCMS method 1; depicted is the extracted ion chromatogram (*m/z* = 387.2) as well as high resolution mass spectrometry and MSMS data corresponding to the formation of (20*S*)-10F-fluorocorynantheidine **(25)** and (20*R*)-10F-fluorocorynantheidine **(26)**; **(b)** *Catharanthus roseus* strictosidine synthase (*Cr*STR), *Catharanthus roseus* strictosidine glucosidase (*Cr*SGD), *Cp*DCS and *Ms*EnolMT were transiently expressed in *Nicotiana benthamiana* and infiltrated with 5F-tryptamine **(21)** and secologanin **(19)**; methanol extracts were analysed using LCMS method 1; depicted is the extracted ion chromatogram (*m/z* = 387.2) as well as high resolution mass spectrometry and MSMS data corresponding to the formation of (20*R*)-10F-fluorocorynantheidine **(26)**.

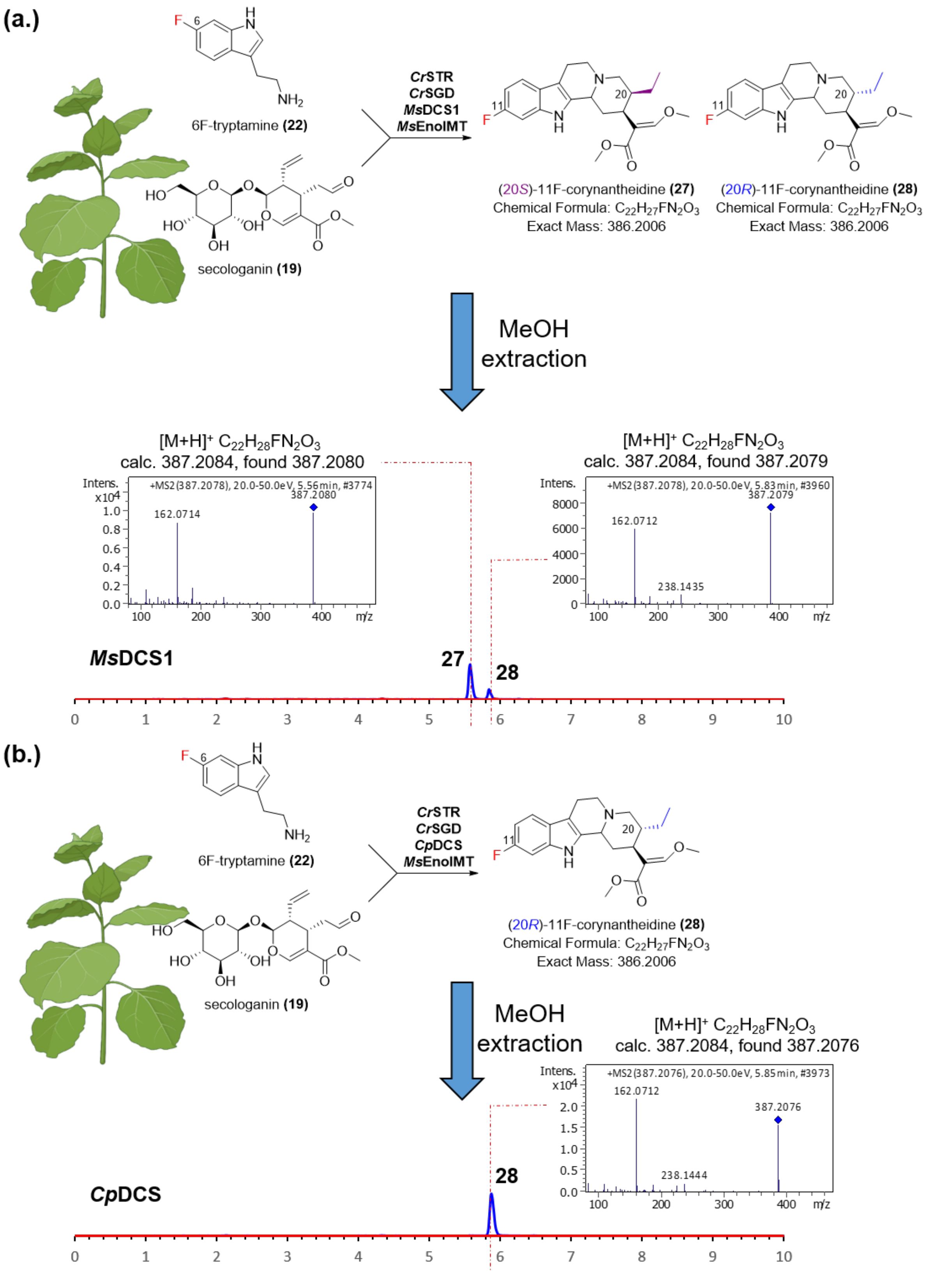


**Supplementary Figure S19 |** See next page for caption.

**Supplementary Figure S19 |** Production of compound corresponding to 11-fluorocorynantheidine. **(a)** *Catharanthus roseus* strictosidine synthase (*CrSTR*), *Catharanthus roseus* strictosidine glucosidase (*CrSGD*), *MsDCS1* and *MsEnolMT* were transiently expressed in *Nicotiana benthamiana* and infiltrated with 6F-tryptamine **(22)** and secologanin (**19**); methanol extracts were analysed using LCMS method 1; depicted is the extracted ion chromatogram (*m/z* = 387.2) as well as high resolution mass spectrometry and MSMS data corresponding to the formation of (20*S*)-11F-fluorocorynantheidine **(27)** and (20*R*)-11F-fluorocorynantheidine **(28)**; **(b)** *Catharanthus roseus* strictosidine synthase (*CrSTR*), *Catharanthus roseus* strictosidine glucosidase (*CrSGD*), *CpDCS* and *MsEnolMT* were transiently expressed in *Nicotiana benthamiana* and infiltrated with 6F-tryptamine **(22)** and secologanin **(19)**; methanol extracts were analysed using LCMS method 1; depicted is the extracted ion chromatogram (*m/z* = 387.2) as well as high resolution mass spectrometry and MSMS data corresponding to the formation of (20*R*)-11F-fluorocorynantheidine **(28)**.

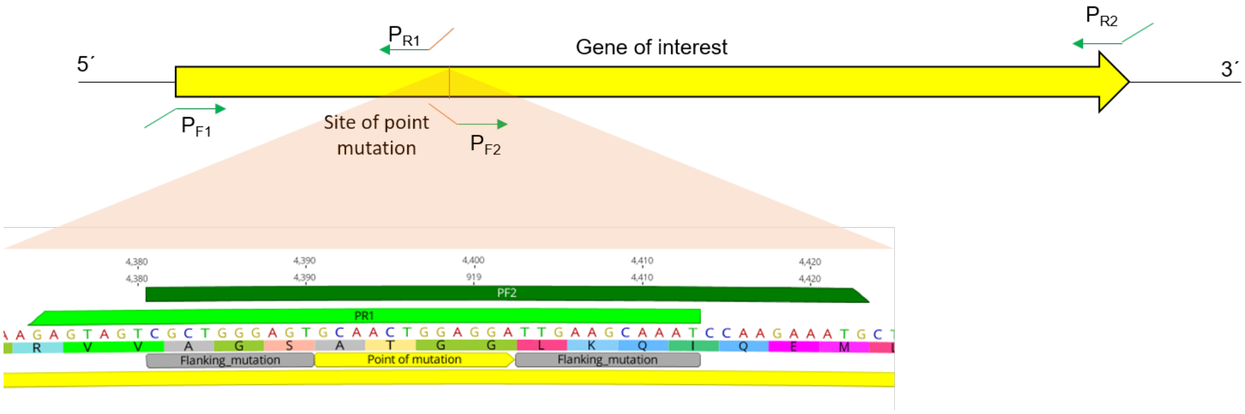


**Supplementary Figure S20 | Cloning strategy to introduce point mutations into selected ADH constructs.**

### Supplementary Tables

**Supplementary Table S1 |** Nucleotide sequences for genes cloned and described in this study; start codons are highlighted in **bold**; stop codons are underlined

| Gene Name | Nucleotide sequence |
|---|---|
| ***Ms*DCS1** | **ATG**GCAGGAAAATGTGCCCAAGAAGAGCACACAGTGAAGGCTTTTGG<br>ATGGGCCGCTAGAGAAGCCTCCGGCGCTCTATCTCCTTACGGGTTCT<br>CAAGAAGGGCAACAGGAGAGCGTGATGTTCGGGTTAAAATTTTGTATT<br>GTGGAATCTGTAGAACAGACGCAGAAATGATCAGCGACAAATTTTGCT<br>TTACTAAGTATCCTCATGTGCCTGGGCATGAGATCGTGGGTGTGGTAT<br>CTGAAGTTGGTAACAAGGTGCAAAAATTCAAGGTTGGAGCTAAAGTCG<br>GTGTGACAGGCATAATTGGATGTTGTCGAACTTGTTATAGCTGTACCA<br>ATGGTCTTGAGAGTTACTGCCCAAATGTTGCACTAACAGAAGCAGGTG<br>AAGGTGGTTGCTCTAACTTCATAGTTTTGGATGAAGACTTTGTGTTTCG<br>TTGGCCTGAGAAATTACCTCTTGATCTTGGAGCTCCTCTCCTGTGTGC<br>TGGAGCCGCTTCTTACAGCCCTTTGAAAAATTTTGGACTTGATAAACC<br>TGGATTGCATATTGGTATAGCTGGTCTTGGTGGCATGGGCCATGTAG<br>CTGTAAAATTTGCTAAGGCTTTTGGGGCAAAGGTGACAGTAATTAGTA<br>CATCAGATAACAAAAAGGAGGAAGCCATTAAAAAATATGGTGCAGACG<br>CATTTTTGAATAGTAGTAATCCTGAGCAGATGCGGGCTGCAGCTGGTA<br>CACTGGCTGCCATCGTTGATACTATCCCTTCGCCTCACTCTCTAGTGC<br>CATTGCTCGATTTATTGTTGCCTCATGGGAAGGTTATTGTATTAGGGG<br>CACCCAGTGAGCCATTTGTGTTGCCGGTTATGCCCCTGCTTCAAGGT<br>GGAAGAGTAGTCGCTGGGAGTTCCGGTGCAAGTTTGAAGCAAATCCA<br>AGAAATGCTCGATTTTGCTGCAGAACACAACATAGTAGCTGATGCTGA<br>GGTTATCCCAATTGACTATATAAACACTGCAATAAAGCGCATTGAGAA<br>GGGCGATATCAAATACCGATTTGTCGTTGACATCGGGAATACACTGAA<br>ATCGGCTTAA |
| ***Ms*DCS2** | **ATG**GCCGAAAAATCACCTGAAGAGGAGCACCCAGTGAAGGCCTTTGG<br>ATTGGCAGCTAAGGACTCATCTGGGATTCTCTCCCCTTTCAACTTCTC<br>AAGAAGGGCAACAGGAGACCATGATGTGCAGCTCAGAGTACTATATT<br>GTGGTCTCTGTTATTATGATACGAAAATGATCAAGAACAAAAGGGGTG<br>TAACTCGCTATCCCTTCGTGTTTGGGCATGAGATTGTGGGTGAAGTAA<br>CTGAGATTGGTAGAGAAGTGCAAAAGTTCAAAGTTGGGGATAAAGTAG<br>GCGTGGGATGCATGGTCGCGTCATGTCGCTCGTGTGAAAGTTGTGCC<br>AACAATTGTGAAAACTACTGCCCAAATGTCTCAGTAACAGATGGGGCA<br>TTTTTCTTCAAGACTGGAGAAGTTCTGTATGGTGGTTGTTCAGACATCA<br>TGGTTGCTGATGAAAATTTTGTGATCCGCTGGCCTGAAAACTTTCCTC<br>TGGATGCTGGGGCTCCTCTCTTGTGTGCTGGGATCACTACTTACAGC<br>CCCTTGAGGAATTTCGGTCTTGATAAACCTGGAATACATGTTGGTATA<br>TATGGTCTTGGTGGACTTGGCCACGTGGCTGTGCAATTTGCCAAGGC<br>TTTTGGGGCAAAAGTGACTGTTATCAGTTCATCTGATAGGAAAAGGAT<br>GGAGGCCATTGAAAAACTTGGTGCAGACTCATTTTTAGTCAACAGTAA<br>TTTGGAGGAAATGCAAGCTGCAATGGGAACAATGCATGGTATCATTGA<br>TACTGTCCCAGCTAATCATTCACTGGTGCCATTGCTTGATTTATTGAAG<br>CCCCAAGGGAAGCTTATCGTTGTAGGTGGACCAGAAAAACCATTTGAA<br>CTACCTGTTTTTCCCCTGCTTCAAGGTGGGAGATTAGTGGCAGGCAGT<br>GCAACTGGAGGAATAAAGCAAACACAAGAAATGATTGATTTTGCAGCT |

| | |
|---|---|
| | GAGCATAACATATTACCACATGTTGAAGTTGTCTCAGTTGATTATGTTA<br>ACACTGCGATAGAGCGCACAGAGAGAGGTGATGTCAAATATAGATTC<br>GTGATTGACATCGGGAATACATTATATTAA |
| ***Ms*EnolMT** | **ATG**CAACCACAGAGAGGGAGAAAGAGAGAGAGAGAGAGAGATAGAG<br>AAGAGATGGAATCCGTGCAGAGCAACAGTAGTTCTTCTGATCAATTCG<br>CAATGAAAGGTGGAGATGACGACTTCAGTTACACAAAGAATTCCACCT<br>GGCAGAGAGATGCAATTCAAGCAACCAAATTTTTCATTCAAGAATCTAT<br>TGCTGAGAAGCTTGACGTCAATAAATTTTGTGGAAAGGCATTTTGCGT<br>TGCTGATTTGGGATGCTCAGTTGGACCTAACACTTTGATAGCAATGCA<br>GAACATTGTTGAAGCTGTGGAGCTTAAATTCAAAAATAGAAAAGGATT<br>CCATTCTCCCACTATCCCTGAATTTCAAGTCTTCTTTAACGATCATACG<br>GTGAATGATTTCAATACCCTCTTTAGATCTCTCCCAACTGGTCACGAC<br>AAGCGCTATTACGGCGTTGGGGTTCCGGGTTCCTTTTACGGTCGATTA<br>TTTCCTTGTGACTCTATTCACATAATGCACACTTCATTTTCTACACCGT<br>TTCTTTCTCAAGTACCAAAAGAGGTGATTGACAAAAATTCAGCTGCGT<br>GGAATAAAGGAAGGATTCATCACAATTATGCTAAAGCAGATGTTTTGA<br>AGGCTTATGAAGCACAACATGCTGAGGATATCGACTGCTTTTTGACGG<br>CTAGAGCTAAAGAACTGGTCCATGGAGGATTATTGATGGATGTGACTT<br>CATTCCGCCCAGATGGGGTCCCTCATACCCATGTCTTGACTAACATAG<br>GGATGGAGGTATTGGGTTATTGCCTCATGGACTTGGCTGGACTTATC<br>GATGAAGAAAACGTGGATTCCTACAACGTTCCAGTTTATCTTCAATCTC<br>CTGAAGAGTTGAAACAAGCTGTTCAACGGAACAAATACTTCAGTATAG<br>AAAAAATGGAGAGCGTGCCTATGATGATAGATTCAGATGTTTCTGCCA<br>AAGCTCAACAATATTCATTGGGAATGAGGGCCGTAATGGGGGACGTG<br>ATTAGAGAGCAATTTGGAGCGGAGATAGTGGATAAACTCTTTGATTTG<br>TTCAAGAAGAAACTTGAAGAGCATCCTAACTTTGCAAAAGGAGTTGTC<br>CTTGACATGTTTGTTCTCCTTAAACGCAATGCAGAGGATTGA |
| ***Cp*DCS** | **ATG**ATGGCCGGAAAATCTCAAGAAGATGGGCAGACGGTAAAGGCTCT<br>AGGATGGGCCGCTAGGGAAGTTTCTGGGGCGATCTCTCCTTTCGATT<br>TCTCAAGAAGGGCCCCAGGAGAGCGCGATGTGCAGGTTAAAATACTA<br>TATTGTGGAATCTGTAGTTTTGACACAGAAATGATCAATAACAAGTTTG<br>GCTTTACCAGATATCCCTTTGTACTCGGGCATGAGATTGTGGGAGTGG<br>TATCTGAAGTTGGTAGAAAGGTGCAAAAATTCAAGATTGGGGATAAAG<br>TTGGTGTAGGAACCATGATTGGATCTTGTCGCACTTGTTATAGCTGCA<br>CTCACAATCTCGAAAATTACTGCCCAAAAGTTACATTAACAGAAGCAA<br>CTTCTGGTGGTTGTTCTAATCTTGTGATAGCAGATGAGGACTTTGTGT<br>TCCATTGGCCGGTGAATTTGCCTCTTGATCTTGGAGCTCCTCTCCTTT<br>GTGCTGGGATTACTGTTTATAGCCCTTTGAAAAATTTTGAACTTGATAA<br>GCCTGGATTGCGTATTGGTGTGGTTGGTCTTGGTGGTATTGGCCATAT<br>AGCTGTAAAATTTGCCAAGGCTTTTGGGGCTAAGGTGACAGTGATTAG<br>TTCATCAGAAAGTAAAAAGGTTGAAGCCATTGAAAAATATGGTGCAGA<br>TTCCTTTTTGGTTAGCAGTGATCCAGGGCAGATGCTGGCAGCTGCCG<br>GAACCTTGGATGGTGTCATTGATACCGTCCCAGCACCTCACTCTATTT<br>TGCCATTCCTTGATTTACTCTTGCCTCGTGGAAAGCTAATTATATTAGG<br>TGCACCAATGGAGCCATTTGTACTGCCAATCTATCCCCTGCTTCAAGG<br>TGGGAGAGTAGTTGCTGGGAGTGCCACTGGAGGATTGAAACAAATCC<br>AAGAAATGCTTCATTTTGCAGCAGAGCACAACATAGTAGCAGATGGCG<br>AGGTTATCCCAATCGACGACATTAACACTGCGATAAAGCGCATTGAGA<br>AAGGCGATGTCAAATATCGATTTGTGGTTGACATTGGCAATACCTTAA<br>AATCTGCTGGTTCGAGAGACCTAGGCTAG |

**Supplementary Table S2 |** Codon-optimized nucleotide sequences for heterologous expression in *Escherichia coli* or *Nicotiana benthamiana*; Start codons are highlighted in **bold**; stop codons are underlined.

| Gene Name | Nucleotide sequence |
|---|---|
| ***Ms*DCS1(*Escherichia coli*)** | **ATG**GCTGGCAAGTGCGCGCAGGAAGAACATACGGTTAAAGCGTTCGG<br>GTGGGCAGCACGAGAGGCGTCAGGTGCGTTAAGTCCGTATGGGTTTT<br>CCCGTCGGGCGACCGGCGAACGCGACGTGCGTGTGAAGATCCTGTA<br>CTGCGGTATTTGCCGCACTGATGCGGAGATGATTTCCGATAAGTTCTG<br>TTTCACGAAATACCCCCACGTTCCAGGTCACGAAATTGTTGGGGTAGT<br>TTCCGAGGTGGGCAATAAAGTTCAGAAGTTTAAAGTAGGCGCGAAGG<br>TGGGAGTCACGGGAATTATAGGTTGCTGCCGCACGTGCTACTCGTGC<br>ACGAACGGCCTGGAATCTTATTGTCCGAACGTGGCCTTGACCGAGGC<br>GGGCGAGGGTGGGTGTAGTAATTTTATTGTACTCGACGAGGATTTCG<br>TATTCCGGTGGCCGGAAAAGCTGCCCCTGGACCTGGGTGCGCCCCT<br>GCTTTGCGCAGGCGCGGCGTCCTATTCACCGCTTAAGAACTTCGGTT<br>TAGACAAGCCGGGCCTGCACATCGGAATCGCGGGCCTGGGTGGAAT<br>GGGACACGTGGCCGTCAAGTTCGCAAAAGCATTCGGCGCCAAAGTAA<br>CCGTTATCTCTACGAGTGACAATAAGAAAGAAGAGGCGATCAAGAAGT<br>ACGGAGCGGATGCTTTCTTAAACTCTTCAAACCCGGAACAAATGCGC<br>GCCGCTGCGGGCACTTTGGCCGCGATTGTGGACACCATTCCAAGTCC<br>GCATTCACTGGTCCCTCTTCTGGACCTTCTTTTACCGCACGGTAAAGT<br>AATCGTTCTGGGCGCGCCTTCCGAACCGTTCGTCTTACCAGTCATGC<br>CGTTATTGCAGGGCGGCCGCGTCGTGGCGGGCAGCAGCGGGGCTTC<br>ACTTAAACAGATTCAGGAGATGTTGGACTTCGCGGCCGAGCATAATAT<br>TGTGGCAGACGCCGAAGTAATTCCGATAGATTACATCAATACAGCGAT<br>CAAACGTATCGAGAAAGGGGACATTAAGTATCGGTTCGTAGTCGATAT<br>TGGCAACACTTTAAAGAGCGCGTGA |
| ***Ms*DCS2 (*Escherichia coli*)** | **ATG**GCAGAGAAGAGTCCCGAGGAAGAACATCCGGTTAAAGCATTCGG<br>GCTCGCGGCCAAAGATAGCTCAGGCATCTTATCTCCGTTTAATTTTAG<br>TCGTCGCGCTACGGGCGATCACGACGTTCAACTGCGTGTTCTGTACT<br>GCGGATTGTGCTACTACGACACTAAGATGATTAAGAATAAGAGAGGC<br>GTGACCAGATACCCGTTTGTATTCGGACACGAAATCGTTGGAGAGGTT<br>ACCGAAATAGGCCGTGAGGTTCAGAAATTTAAGGTAGGTGACAAGGT<br>TGGAGTCGGTTGTATGGTTGCTTCCTGCCGTAGTTGCGAGTCTTGCG<br>CGAATAACTGCGAGAATTATTGTCCGAACGTATCCGTGACGGACGGT<br>GCGTTCTTCTTTAAAACGGGTGAGGTGTTGTACGGCGGCTGCTCCGA<br>TATTATGGTCGCGGACGAGAACTTCGTTATTCGGTGGCCAGAGAATTT<br>CCCACTTGACGCAGGAGCGCCATTACTGTGCGCAGGCATTACAACGT<br>ATTCGCCATTACGCAACTTTGGCCTGGACAAGCCGGGCATCCACGTG<br>GGCATCTACGGACTCGGTGGTCTCGGTCATGTTGCGGTTCAGTTCGC<br>GAAAGCCTTCGGCGCGAAGGTTACAGTCATATCTTCTAGTGACCGGA<br>AGCGGATGGAAGCGATCGAGAAGCTGGGCGCTGATAGCTTCCTGGT<br>GAATAGCAACTTAGAAGAGATGCAGGCAGCTATGGGGACCATGCACG<br>GAATTATCGACACCGTTCCGGCGAACCACAGTTTAGTCCCGTTACTGG<br>ACCTGTTAAAACCACAGGGTAAACTGATAGTGGTTGGTGGTCCTGAGA<br>AGCCGTTCGAGCTGCCAGTATTCCCGTTACTGCAGGGTGGCCGTCTG<br>GTTGCTGGAAGCGCCACAGGTGGCATTAAACAGACGCAGGAGATGAT<br>AGACTTCGCCGCGGAACACAATATTCTGCCCCACGTCGAGGTGGTTA<br>GCGTCGACTACGTCAATACGGCAATTGAACGAACGGAACGTGGAGAC<br>GTTAAGTACCGTTTTGTAATCGATATTGGCAACACCCTGTACTGA |

| | |
|---|---|
| ***Cp*DCS** (***Escherichia coli***) | **ATG**GCAGGTAAGAGCCAGGAAGACGGCCAAACAGTTAAAGCCTTAGG<br>GTGGGCGGCACGCGAGGTGTCAGGAGCAATATCGCCGTTTGACTTTT<br>CGCGTAGAGCACCCGGCGAAAGAGACGTTCAAGTGAAGATTCTGTAC<br>TGCGGCATATGCTCATTCGATACCGAGATGATTAACAATAAATTCGGA<br>TTCACTCGCTACCCATTCGTTTTGGGCCACGAAATCGTCGGTGTTGTG<br>TCGGAAGTGGGCCGTAAAGTCCAGAAGTTTAAAATAGGTGACAAGGT<br>AGGCGTGGGCACAATGATCGGGAGCTGCCGTACATGCTACTCATGTA<br>CACATAACTTGGAGAACTATTGTCCGAAGGTGACGCTGACTGAGGCT<br>ACATCCGGCGGCTGCTCCAACCTGGTAATTGCGGACGAGGATTTCGT<br>ATTTCACTGGCCTGTAAACCTGCCGCTGGACCTGGGCGCACCGCTTC<br>TCTGCGCAGGCATAACCGTATACAGTCCATTAAAGAACTTCGAGTTGG<br>ACAAACCCGGCCTTCGCATCGGCGTAGTCGGGCTCGGCGGGATAGG<br>TCACATTGCAGTGAAGTTCGCAAAAGCATTCGGTGCCAAAGTTACCGT<br>TATCTCGTCTTCGGAGTCAAAGAAAGTCGAGGCTATCGAGAAGTACG<br>GCGCCGACTCTTTCCTGGTGAGTTCCGACCCTGGCCAAATGCTTGCT<br>GCGGCGGGGACGTTAGACGGCGTGATCGACACTGTACCTGCCCCAC<br>ATTCCATCTTACCTTTTCTGGACCTTTTATTACCACGGGGCAAACTGAT<br>CATCCTGGGCGCGCCCATGGAACCGTTCGTTCTCCCTATTTACCCGC<br>TTCTCCAGGGTGGCCGTGTGGTAGCAGGATCGGCGACAGGTGGGCT<br>TAAGCAGATACAGGAGATGTTGCACTTCGCTGCCGAACATAATATTGT<br>TGCGGACGGTGAAGTAATTCCCATAGATGATATAAATACGGCAATTAA<br>ACGTATCGAGAAAGGGGACGTTAAGTACCGGTTCGTTGTGGATATCG<br>GTAACACATTGAAGAGTGCATAA |
| **PsiH** (***Nicotiana benthamiana***) | **ATG**ATTGCAGTTTTATTTTCATTCGTGATCGCCGGATGTATTTACTACA<br>TCGTTTCACGGAGAGTTAGAAGAAGTAGGTTGCCACCGGGTCCACCC<br>GGGATACCTATTCCATTCATAGGAAACATGTTCGACATGCCTGAAGAA<br>TCTCCGTGGTTGACTTTCCTCCAGTGGGGTAGAGACTACAACACAGAT<br>ATCTTGTACGTAGATGCAGGTGGAACTGAGATGGTGATATTAAATACT<br>CTTGAGACAATCACTGATCTGTTGGAGAAAAGAGGCTCTATTTACTCT<br>GGAAGACTAGAATCTACTATGGTTAATGAACTTATGGGTTGGGAATTT<br>GACTTAGGTTTTATCACATATGGCGATAGATGGAGAGAGGAGAGGCG<br>TATGTTCGCCAAGGAATTTAGTGAGAAGGGTATAAAGCAATTTAGGCA<br>TGCACAGGTGAAAGCTGCGCATCAACTTGTACAACAATTAACAAAGAC<br>CCCAGATAGGTGGGCACAACATATTCGACATCAAATCGCGGCAATGA<br>GTTTAGACATTGGCTATGGAATTGATCTCGCAGAGGATGATCCTTGGT<br>TAGAGGCTACACATCTCGCTAATGAGGGTCTAGCTATTGCGTCCGTAC<br>CTGGCAAATTTTGGGTCGATAGTTTTCCTTCTTTAAAGTACCTTCCTGC<br>TTGGTTTCCAGGGGCAGTTTTTAAGCGCAAGGCCAAAGTCTGGAGAG<br>AAGCTGCTGACCATATGGTAGACATGCCATACGAGACAATGCGGAAA<br>CTAGCTCCTCAAGGTCTTACTCGACCATCCTACGCGTCAGCGAGGTTA<br>CAAGCTATGGATTTGAATGGCGACCTTGAACATCAAGAACACGTCATC<br>AAAAATACTGCTGCTGAAGTAAATGTCGGGGGAGGGGATACAACTGT<br>CTCTGCAATGTCTGCTTTCATTCTCGCAATGGTGAAGTATCCTGAAGT<br>ACAACGTAAGGTTCAGGCTGAATTGGACGCATTGACAAATAATGGCCA<br>AATTCCAGATTACGACGAAGAAGATGATTCTCTTCCTTACCTAACCGC<br>TTGTATTAAAGAGTTATTCAGATGGAACCAAATTGCGCCCTTAGCTATT<br>CCTCATAAGTTGATGAAAGATGATGTATATCGAGGATATCTGATTCCTA<br>AGAACACATTAGTTTTCGCTAATACATGGGCCGTTCTAAACGATCCTG<br>AGGTGTATCCTGACCCAAGTGTGTTCCGGCCAGAGCGTTACCTGGGA<br>CCAGACGGGAAGCCAGATAATACCGTTCGAGATCCTCGTAAAGCAGC<br>TTTTGGGTATGGTCGGAGGAATTGTCCAGGCATTCATCTAGCTCAATC |

| | |
|---|---|
| | AACAGTGTGGATTGCCGGAGCTACTTTGCTTTCTGCTTTCAACATCGA GCGACCTGTTGATCAAAATGGCAAGCCCATCGACATTCCTGCTGATTT CACAACGGGGTTCTTTAGACATCCTGTGCCATTCCAATGTCGTTTTGT GCCCAGGACAGAACAGGTGTCACAGTCAGTTAGTGGCCCTTGA |

**Supplementary Table S3 |** List of expression plasmids generated in this study; all plasmids were generated by In-Fusion cloning, as described above; oligonucleotides used for the construction of each plasmid are listed in Supplementary Table S4.

| # | Plasmid | Template DNA for PCR | Primers used for construction |
|---|---|---|---|
| *Plasmids constructed for the purpose of transient gene expression in Nicotiana benthamiana* | | | |
| **P1** | 3Ω1_MsDCS1 | cDNA kratom | #1, #2 |
| **P2** | 3Ω1_MsDCS2 | cDNA kratom | #3, #4 |
| **P3** | 3Ω1_CpDCS | Synthetic gene | #5, #6 |
| **P4** | 3Ω1_PsiH | Synthetic gene | #7, #8 |
| **P24** | 3Ω1_MsEnolMT | cDNA kratom | #37, #38 |
| *Plasmid containing point mutations to probe the stereoselectivity of MsDCS1 and CpDCS destined for transient gene expression in Nicotiana benthamiana* | | | |
| **P5** | 3Ω1_MsDCS1_S295A_G296T_A297G_S298G | P1 | #9, #10, #11, #12 |
| **P6** | 3Ω1_MsDCS1_I100M | P1 | #9, #12, #13, #14 |
| **P7** | 3Ω1_MsDCS1_S116N | P1 | #9, #12, #15, #16 |
| **P8** | 3Ω1_MsDCS1_S295A_G296T_A297G_S298G_I100M | P5 | #9, #12, #13, #14 |
| **P9** | 3Ω1_MsDCS1_S295A_G296T_A297G_S298G_S116N | P5 | #9, #12, #15, #16 |
| **P10** | 3Ω1_MsDCS1_S295A_G296T_A297G_S298G_I100M_S116N | P8 | #9, #12, #15, #16 |
| **P11** | 3Ω1_MsDCS1_T53F | P1 | #9, #12, #17, #18 |
| **P12** | 3Ω1_MsDCS1_S295A_G296T_A297G_S298G_I100M_S116N_T53F | P10 | #9, #12, #17, #18 |
| **P13** | 3Ω1_CpDCS_A295S_T296G_G297A_G298S | P3 | #5, #6, #19, #20 |
| **P14** | 3Ω1_CpDCS_M100I | P3 | #5, #6, #21, #22 |
| **P15** | 3Ω1_CpDCS_N116S | P3 | #5, #6, #23, #24 |
| **P16** | 3Ω1_CpDCS_A295S_T296G_G297A_G298S_M100I | P13 | #5, #6, #21, #22 |
| **P17** | 3Ω1_CpDCS_A295S_T296G_G297A_G298S_N116S | P13 | #5, #6, #23, #24 |
| **P18** | 3Ω1_CpDCS_A295S_T296G_G297A_G298S_M100I_N116S | P16 | #5, #6, #23, #24 |
| **P19** | 3Ω1_CpDCS_A295S_T296G_G297A_G298S_M100I_N116S_F53T | P18 | #5, #6, #25, #26 |
| *Plasmids constructed for the bacterial expression of $His_6$-tagged genes* | | | |
| **P20** | pOPINF-$His_6$-MsDCS1 | Synthetic gene | #27, #28 |
| **P21** | pOPINF-$His_6$-MsDCS2 | Synthetic gene | #29, #30 |
| **P22** | pOPINF-$His_6$-CpDCS | Synthetic gene | #31, #32 |
| **P23** | pOPINM-$His_6$-MsEnolMT | cDNA kratom | #35, #36 |

**Table S4 |** Oligonucleotides used for the construction of 3Ω1/pOPINM expression plasmids destined for transient gene expression in *Nicotiana benthamiana* or *E. coli.*

| **Oligonucleotide** | |
|---|---|
| *Oligonucleotides used for subcloning of wild-type gene sequences* | |
| #1 | TTTATGAATTTTGCAGCTCG<br>ATGGCAGGAAAATGTGCCCAAG |
| #2 | GACAACCACAACAAGCACCGT<br>TAAGCCGATTTCAGTGTATTCCCGATG |
| #3 | TTTATGAATTTTGCAGCTCG<br>ATGGCCGAAAAATCACCTGAAGAGGAG |
| #4 | GACAACCACAACAAGCACCGT<br>CATATCCAACCAAAAAATTGAAACAAAGGAAATG |
| #5 | TTTATGAATTTTGCAGCTCG<br>ATGATGGCCGGAAAATCTCAAG |
| #6 | GACAACCACAACAAGCACCG<br>CTAGCCTAGGTCTCTCG |
| #7 | TTTATGAATTTTGCAGCTCG<br>ATGATTGCAGTTTTATTTTCATTCGTGATCG |
| #8 | GACAACCACAACAAGCACCGT<br>CAAGGGCCACTAACTGACTGTGAC |
| *Oligonucleotides used to introduce point mutations into MsDCS1 or CpDCS* | |
| #9 | TTTATGAATTTTGCAGCTCGATGGCAGGAAAATGTGCCC |
| #10 | ATTTGCTTCAATCCTCCAGTTGCACTCCCAGCGACTACTC |
| #11 | CGCTGGGAGTGCAACTGGAGGATTGAAGCAAATCCAAGAAATG |
| #12 | GACAACCACAACAAGCACCGTTAAGCCGATTTCAGTGTATTCC |
| #13 | ACAACATCCAATCATGCCTGTCACACCG |
| #14 | CGGTGTGACAGGCATGATTGGATGTTGTC |
| #15 | CATTTGGGCAGTAATTCTCAAGACCATTG |
| #16 | CAATGGTCTTGAGAATTACTGCCCAAATG |
| #17 | CATTTCTGCGTCGAATCTACAGATTCCAC |
| #18 | GTGGAATCTGTAGATTCGACGCAGAAATGATC |
| #19 | GATTTGTTTCAAACTTGCACCGGAACTCCCAGCAAC |
| #20 | GTTGCTGGGAGTTCCGGTGCAAGTTTGAAACAAATC |
| #21 | GACAAGATCCAATTATGGTTCCTACACCAAC |
| #22 | GTTGGTGTAGGAACCATAATTGGATCTTGTC |
| #23 | GTAACTTTTGGGCAGTAACTTTCGAGATTGTGAGTG |
| #24 | CACTCACAATCTCGAAAGTTACTGCCCAAAAGTTAC |
| #25 | CATTTCTGTGTCTGTACTACAGATTCCAC |
| #26 | GTGGAATCTGTAGTACAGACACAGAAATG |
| #27 | AAGTTCTGTTTCAGGGCCCGATGGCTGGCAAGTGCGC |
| #28 | ATGGTCTAGAAAGCTTTATCACGCGCTCTTTAAAGTGTTG |
| #29 | AAGTTCTGTTTCAGGGCCCGATGGCAGAGAAGAGTCCCG |
| #30 | ATGGTCTAGAAAGCTTTATCAGTACAGGGTGTTGCCAATAT |
| #31 | AAGTTCTGTTTCAGGGCCCGATGGCAGGTAAGAGCCAGG |
| #32 | ATGGTCTAGAAAGCTTTATTATGCACTCTTCAATGTGTTACC |
| #33 | TGTCGCTTTGGTTCTCAAGG |
| #34 | TGCTCAGAACTGCAATCAGA |
| #35 | AAGTTCTGTTTCAGGGCCCGATGCAACCACAGAGAGGGAGAAAGAG |
| #36 | ATGGTCTAGAAAGCTTTAATCCTCTGCATTGCGTTTAAGGAGAACAAAC |
| #37 | TTTATGAATTTTGCAGCTCGATGGAATCCGTGCAGAGCAACAGTAGTTC |
| #38 | GACAACCACAACAAGCACCGTCAATCCTCTGCATTGCGTTTAAGGAGAACAAAC |
